# SNM1A (*DCLRE1A*) is crucial for efficient repair of complex DNA breaks in human cells

**DOI:** 10.1101/2022.07.21.500940

**Authors:** Lonnie P. Swift, B. Christoffer Lagerholm, Hannah T. Baddock, Lucy R. Henderson, Malitha Ratnaweera, Blanka Sengerova, Sook Lee, Abimael Cruz-Migoni, Dominic Waithe, Christian Renz, Helle D. Ulrich, Joseph A. Newman, Christopher J. Schofield, Peter J. McHugh

**Affiliations:** Department of Oncology, MRC-Weatherall Institute of Molecular Medicine, University of Oxford, John Radcliffe Hospital, Oxford OX3 9DS, UK; Wolfson Imaging Centre, MRC-Weatherall Institute of Molecular Medicine, University of Oxford, John Radcliffe Hospital, Oxford OX3 9DS, UK; Institute of Molecular Biology gGmbH (IMB), Ackermannweg 4, 55128 Mainz, German; Centre for Medicines Discovery, University of Oxford, Oxford OX3 7DQ, United Kingdom; Chemistry Research Laboratory, Department of Chemistry and the Ineos Oxford Institute for Antimicrobial Research, University of Oxford, Oxford, OX1 3TA, UK; Kennedy Institute of Rheumatology, University of Oxford, Roosevelt Drive, Oxford, OX3 7FY, UK; Calico Life Sciences, 1170 Veterans Blvd, South San Francisco, CA, 94080, USA; Institute of Organic Chemistry and Biochemistry of the Czech Academy of Sciences. Flemingovo náměstí 542/2 160 00 Praha 6. Czech Republic

## Abstract

DNA double-strand breaks (DSBs), such as those produced by radiation and radiomimetics, are amongst the most toxic forms of cellular damage, in part because they involve extensive oxidative modifications at the break termini. Prior to completion of DSB repair, the chemically modified termini must be removed. Various DNA processing enzymes have been implicated in the processing of these ‘dirty ends’, but molecular knowledge of this process is limited. Here, we demonstrate a role for the metallo-β-lactamase fold 5ʹ-3ʹ exonuclease SNM1A in this vital process. Cells disrupted for SNM1A manifest increased sensitivity to radiation and radiomimetic agents and show defects in DSB damage repair. SNM1A is recruited and is retained at the sites of DSB damage *via* the concerted action of its three highly conserved PBZ, PIP box and UBZ interaction domains, which mediate interactions with poly-ADP-ribose chains, PCNA and the ubiquitinated form of PCNA, respectively. SNM1A can resect DNA containing oxidative lesions induced by radiation damage at break termini. The combined results reveal a crucial role for SNM1A to digest chemically modified DNA during the repair of DSBs and imply that the catalytic domain of SNM1A is an attractive target for potentiation of radiotherapy.

## Introduction

DNA double-strand breaks (DSBs) are one of the most cytotoxic forms of damage, with even small number of unrepaired breaks being potentially lethal^1^. In human cells, the majority of DSBs are resolved by one of two pathways, that is non-homologous end-joining (active from G1- through to the G2-phase of the cell cycle) or homologous recombination (which is activated as cells traverse S-phase)^2^. Prior to completion of DSB repair, any chemically modified nucleotides or aberrant structures must be removed from the break-ends. This process is especially important for DSBs induced by ionising radiation (IR) or radiomimetic drugs, including bleomycin and related agents, which are associated with extensive, mostly oxidative, DNA modifications at the break termini^3, 4^. Various factors have been implicated in processing such ‘dirty end’ ends, including tyrosyl DNA phosphodiesterase (TDP1)^5^, polynucleotide kinase (PNK)^6^, aprataxin^7–10^ and Artemis/SNM1C (discussed below)^11, 12^. TDP1 can remove the 3ʹ-phosphoglycolate ends that constitute approximately 10% of the termini produced by IR^13^. PNK catalyses removal of 3ʹ-phosphate groups and addition of phosphates to 5ʹ-hydroxyl moieties in preparation for end ligation^6^, modifications that are associated with oxidation reactions following IR. Aprataxin catalyses deadenylation releasing DNA Ligase IV during abortive ligation reactions during NHEJ, removing the associated AMP group^14^.

The SNM1A 5ʹ-3ʹ exonuclease encoded by the *DCLRE1A* gene is a member of a family of metallo-β-lactamase (MBL) fold DNA nucleases conserved, at least, from yeasts to humans^15^. The family is characterised presence of a β-CASP (CPSF-Artemis-SNM1A-Pso2) domain that together with MBL domain forms the nuclease active site^16^. Human cells have three MBL-β-CASP DNA nuclease paralogues: SNM1A, SNM1B (also known as Apollo) and SNM1C (Artemis)^16^.

SNM1A has a key role in DNA interstrand crosslink repair (ICL repair), where a role in the replication-coupled pathway of ICL repair in mammalian cells is mediated through interaction of SNM1A with PCNA *via* its PCNA interacting protein (PIP) motif as well as interaction *via* an N-terminally located ubiquitin-binding zinc finger (UBZ) with the monoubiquitinated form of PCNA^15, 17, 18^. Very recently, SNM1A has been shown to be important in mediating DNA damage tolerance associated with break-induced replication and recombination at telomeres maintained by the alternative lengthening (ALT) pathway^19^. While the role of SNM1B/Apollo in DNA repair has not been comprehensively elucidated, although it has been implicated in DSB and ICL repair, it has a well-defined role in processing of newly-synthesised telomere leading-strands to maintain the structure at telomere termini^20^.

SNM1A and SNM1B/Apollo are both 5ʹ-3ʹ exonucleases^21^. SNM1C/Artemis has a distinct catalytic profile, with limited 5ʹ-3ʹ exonuclease activity. On association with DNA-PKcs (the catalytic subunit of the NHEJ factor DNA-PK), the 5ʹ-3ʹ endonuclease activity of SNM1C/Artemis acquires capacity to open DNA hairpins and to remove overhangs and unpaired regions at damaged / aberrant DNA termini^12, 22^. SNM1C/Artemis plays a key role in V(D)J recombination, by removing RAG-generated hairpin intermediates and in processing of DNA breaks bearing modified termini, including oxidised nucleotides induced by IR as part of the NHEJ pathway

Here we report that cell lines disrupted for SNM1A by genome-editing display the anticipated increased sensitivity to DNA crosslinking agents. Unexpectedly, screening more generally for DNA damage sensitivity revealed the sensitivity of SNM1A disrupted cells to radiation and radiomimetic agents. This unanticipated observation led us to comprehensively characterise a DSB repair role for SNM1A. We demonstrate that several ligand associations act to recruit SNM1A to DSBs and that its DSB repair role likely reflects the exceptional capacity of SNM1A to hydrolyse DNA break substrates with chemical and structural modifications.

## Results

### SNM1A^-^ cells show a profound sensitivity to radiometric damage

To investigate the phenotype of human cells disrupted for SNM1A we utilised both zinc-finger Nuclease (ZFN)^23^ and CRISPR-Cas9^24^ gene editing to generate SNM1A^-^ cells. Initially, we employed ZFN constructs targeting the first exon of SNM1A. A putative U2OS SNM1A disrupted clone was identified, containing a 22-nucleotide deletion in exon 1, producing a frameshift and premature stop codon distal from the deletion site (Suppl. Fig. 1); SNM1A protein was not detected by immunoblotting in these cells (Fig. 1A).

**Figure 1.**
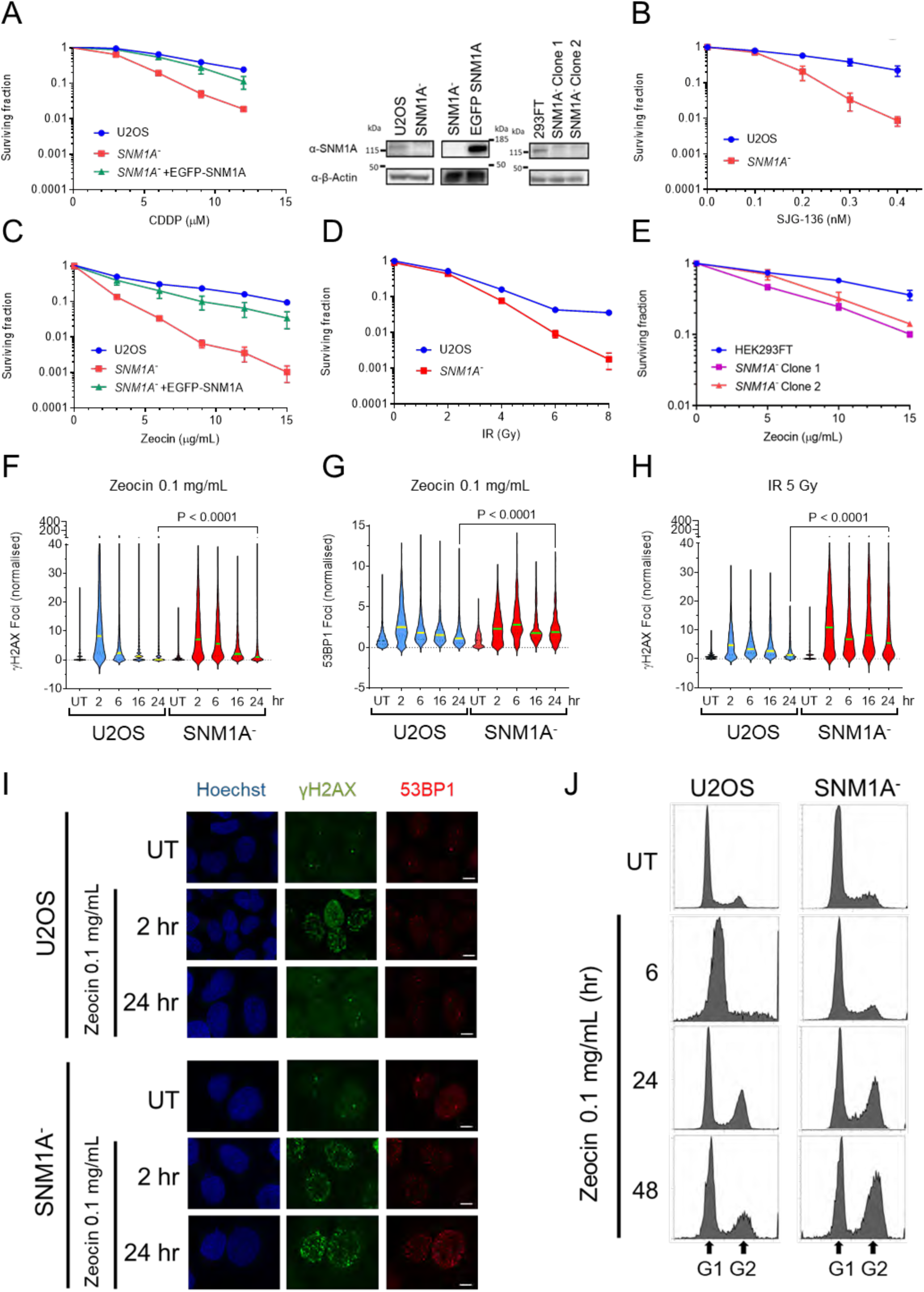
SNM1A^-^ cells show a profound sensitivity to radiometric damage. **A** SNM1A^-^ cells generated in U2OS background are sensitive to Cisplatin (CDDP) treatment in a clonogenic survival assay and the sensitivity is suppressed in cells stably expressing an EGFP-SNM1A (left hand panel). SNM1A protein is not detected in the SNM1A^-^ cells by immunoblot (right hand panel), compared to wildtype U2OS cells, and the EGFP-SNM1A band is evident in the complemented cell line by immunoblot analysis as shown in the inset panels. **B.** SNM1A^-^ cells are also sensitive to the DNA crosslinking agent SJG-136. **C.** SNM1A^-^ cells show a profound sensitivity to the radiomimetic Zeocin and this sensitivity is rescued by expressing EGFP-SNM1A. **D.** SNM1A^-^ cells are also sensitive to ionising radiation. **E.** Zeocin sensitivity is also observed in SNM1A^-^ 293FT cells, SNM1A protein is not detected in these cell lines by immunoblot analysis (as shown in right hand panel **A**). Zeocin treatment induces persistent γH2AX **F** and 53BP1 **G** foci in SNM1A^-^ cells thus indicating a failure to repair Zeocin-induced complex DSB damage. **H.** shows a similar response of γH2AX following IR treatment (for foci data n=3 biological repeats counting >500 cells per repeat, error = SEM). Panel **I.** shows representative foci images of the data from **F. and G.** (Scale bars = 10 µm). **J.** Cell cycle distribution by DNA content determined by flow cytometry following Zeocin treatment over 48 hours reveals a G2 accumulation in SNM1A^-^ cells not observed WT U2OS cells.

To examine the DNA damage response defects in cells lacking SNM1A, we treated them with a broad range of genotoxic agents and determined their survival relative to the parental U2OS cells in clonogenic survival assays. As anticipated from previous work^17^, SNM1A^-^ cells were more sensitive to the crosslinking agents cisplatin and SJG-136, supporting an important role for SNM1A in removing ICLs (Fig. 1A and 1B). Introduction of EGFP-SNM1A into the SNM1A^-^ cell line restored the cisplatin sensitivity of the SNM1A^-^ cells to near wildtype levels (Fig. 1A). Sensitivity to UVC irradiation was observed, possibly as a consequence of the rare ICLs induced by this form of damage (Suppl. Fig. 2A). An absence of increased sensitivity was observed for a range of other genotoxins in SNM1A^-^ cells, including formaldehyde (HCHO), hydrogen peroxide (H2O2) and the topoisomerase 1 inhibitor Campothecin, relative to the parental U2OS cells (Suppl. Fig. 2B, C, D). Strikingly, with the radiomimetic drug Zeocin, dramatically increased sensitisation was observed for the SNM1A^-^ cells. This sensitivity was suppressed by stable complementation with EGFP-SNM1A (Fig. 1C). Since the principal genotoxic lesion induced by Zeocin is DSBs associated with chemically modified termini^25^, we assessed the sensitivity of SNM1A^-^ cells to ionising radiation (IR) which induces related damage (Fig. 1D). Sensitivity to IR was observed, although the overall increase in sensitisation was lower than for Zeocin.

Together the above-described results suggest a role for SNM1A in repair of the major cytotoxic lesions induced by Zeocin and IR, DSBs associated with chemically modified termini. To strengthen this proposal, we used CRISPR-Cas9 to target a sequence in the first exon of SNM1A in 293FT cells (Suppl. Fig. 1) and identified clones with frameshift mutations resulting in a distal introduction of a premature stop codon; these cells do not express SNM1A as determined by immunoblot (Fig. 1A). As with the U2OS SNM1A^-^ cell results, the 293FT cells manifest clearly increased sensitivity to Zeocin treatment (Fig. 1E).

### SNM1A^-^ cells are defective in the repair of radiation and radiomimetic damage

As SNM1A is a 5ʹ-3ʹ repair exonuclease, a plausible explanation for the sensitivity of SNM1A cells to Zeocin and IR is defective processing of the toxic DSBs these agents induce. To determine whether SNM1A^-^ cells exhibit the hallmarks of a DSB repair defect, we initially employed pulsed field gel electrophoresis (PFGE) with the aim of directly detecting Zeocin and IR induced DSBs and monitoring their resolution. However, in the cell lines employed we found that the doses of Zeocin or IR required to permit detection of DSBs were too high for repair to be monitored by PFGE in parental U2OS or 293FT cells. Consequently, we moved to monitoring the dynamics of γH2AX and 53BP1 foci, which are markers of DSB induction^26, 27^. Following treatment of U2OS cells, with Zeocin or IR, γH2AX and 53BP1 levels peaked within 2 hours, and were largely resolved within 24 hours (Fig. 1F, 1G and 1H, representative images for Zeocin foci shown in 1I). However, in SNM1A^-^ cells treated with Zeocin, a substantial fraction of γH2AX and 53BP1 foci persisted after 24 hours (Fig. 1F and 1G). Employing IR as the DSB-inducing agent also led to a marked delay in γH2AX foci resolution in SNM1A cells (Fig. 1H). Analysis of cell cycle progression following Zeocin treatment (0.1 mg/mL) demonstrated that parental (U2OS) cells only transiently slowed and arrested, whereas the SNM1A^-^ cells accumulated in the late S/G2 phase of the cell cycle for up to 48 hours, characteristic of cells accumulating unrepaired DSBs (Fig. 1J, Suppl. Fig. 3). Together, these observations demonstrate that SNM1A-deficient cells are proficient in signalling the presence of damage, but impaired in their repair-response to Zeocin-induced DSBs.

Interestingly, when reporter assays were used to investigate overt defects in NHEJ or HR induced by the ʹcleanʹ DSBs produced by the I-SceI endonuclease, no major repair defects were observed in SNM1A^-^ cells (Suppl. Fig. 4A, B). By contrast, and as anticipated, cells disrupted for XRCC4 or cells depleted for BRCA2, acted as positive controls for defects in NHEJ or HR respectively^28, 29^. A slightly elevated level of HR was observed for SNM1A^-^ cells, implying a minor, but not essential, role in DSB repair pathway utilisation at clean breaks. The lack of an HR defect also suggests that SNM1A does not play a role in the ‘canonical’ pathways of end-resection that are required to generate the requisite 3ʹ-overhangs for HR. Therefore, only DNA DSBs associated with complex, chemically altered termini, such as those induced by radiation and radiomimetics, appear to rely on SNM1A for repair.

We then examined whether SNM1A is recruited to sites of Zeocin-induced DSBs, by employing cells that stably express a functional N-terminally EGFP-tagged form of SNM1A (Fig. 1A). By employing 53BP1 as a marker for DSB induction and localisation, we observed that Zeocin treatment causes the majority of 53BP1 foci to colocalise with EGFP-SNM1A foci (Fig. 2A). High resolution imaging reveals that SNM1A localises immediately adjacent to 53BP1 foci. (Fig. 2B). The observation that SNM1A localises to the vicinity of complex DSBs further supports the proposal that SNM1A plays an important role in responding to complex DSBs.

**Figure 2.**
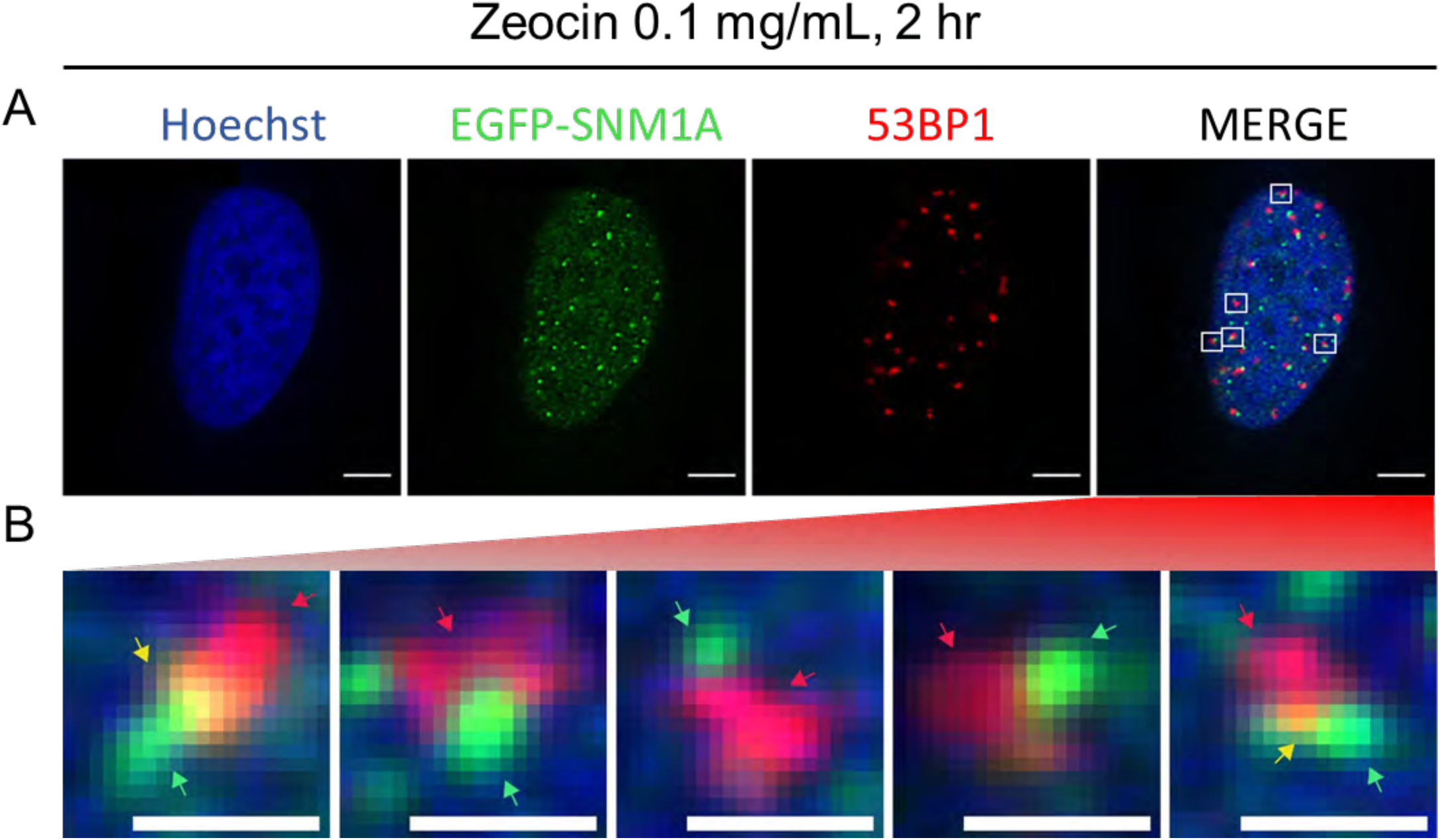
Formation and proximity of SNM1A and 53BP1 foci in response to Zeocin induced DNA damage. **A.** EGFP-SNM1A (green) and 53BP1 (red) foci co-localise at sites of Zeocin-induced DNA damage. EGFP-SNM1A expressing cells treated with Zeocin were fixed and stained with anti-53BP1 (scale bars = 10 µm). **B.** Magnified views of select regions of interest of the co-localised EGFP- SNM1A and 53BP1 foci (yellow) showing the proximity of these foci (scale bars = 1 µm).

### The PBZ, UBZ and PIP box act in concert to recruit SNM1A to complex DNA breaks

The N-terminus of SNM1A is predicted to contain three highly-conserved motifs: a putative poly-ADP-ribose (PAR)-binding zinc finger (PBZ)^15, 30^, a ubiquitin-binding zinc finger (a UBZ4 motif, hereafter UBZ)^15, 31^ which is involved in the localisation of SNM1A to ICLs in S-phase through mediating interaction with ubiquitinated PCNA^18^, and a PIP (PCNA interacting peptide) box^15, 32^ (Fig. 3A; sequence alignments are in Suppl. Fig. 5). Evidence for the direct interaction of these motifs with any of their predicted ligands has not been reported. As structural insight into the non-catalytic N-terminal region of SNM1A is currently lacking, likely due to extensive overall disorder, we predicted the architectures of the UBZ and PBZ domains, and PIP box using the AlphaFold platform^33^. The predicted SNM1A UBZ domain structure is similar to that of the UBZ4 domain of RAD18^34^, except for a change in the conformation of the secondary structure of the Zn^2+^ coordinating residues of its C3H1-type zinc finger (Fig 3B, left-hand panel); the predicted SNM1A PBZ domain structure closely matched the experimental PBZ structure of APLF^35^ with the PBZ residues C155 and C161 being predicted to coordinate Zn^2+^ (Fig 3B, middle panel); the PIP box prediction for SNM1A agreed strongly with the experimental (high affinity) PIP box structure of P21/WAF1^36^, with critical residues Q556, I559, Y562 and F563 lying close to the expected grooves in PCNA (Fig 3B, right-hand panel).

**Figure 3.**
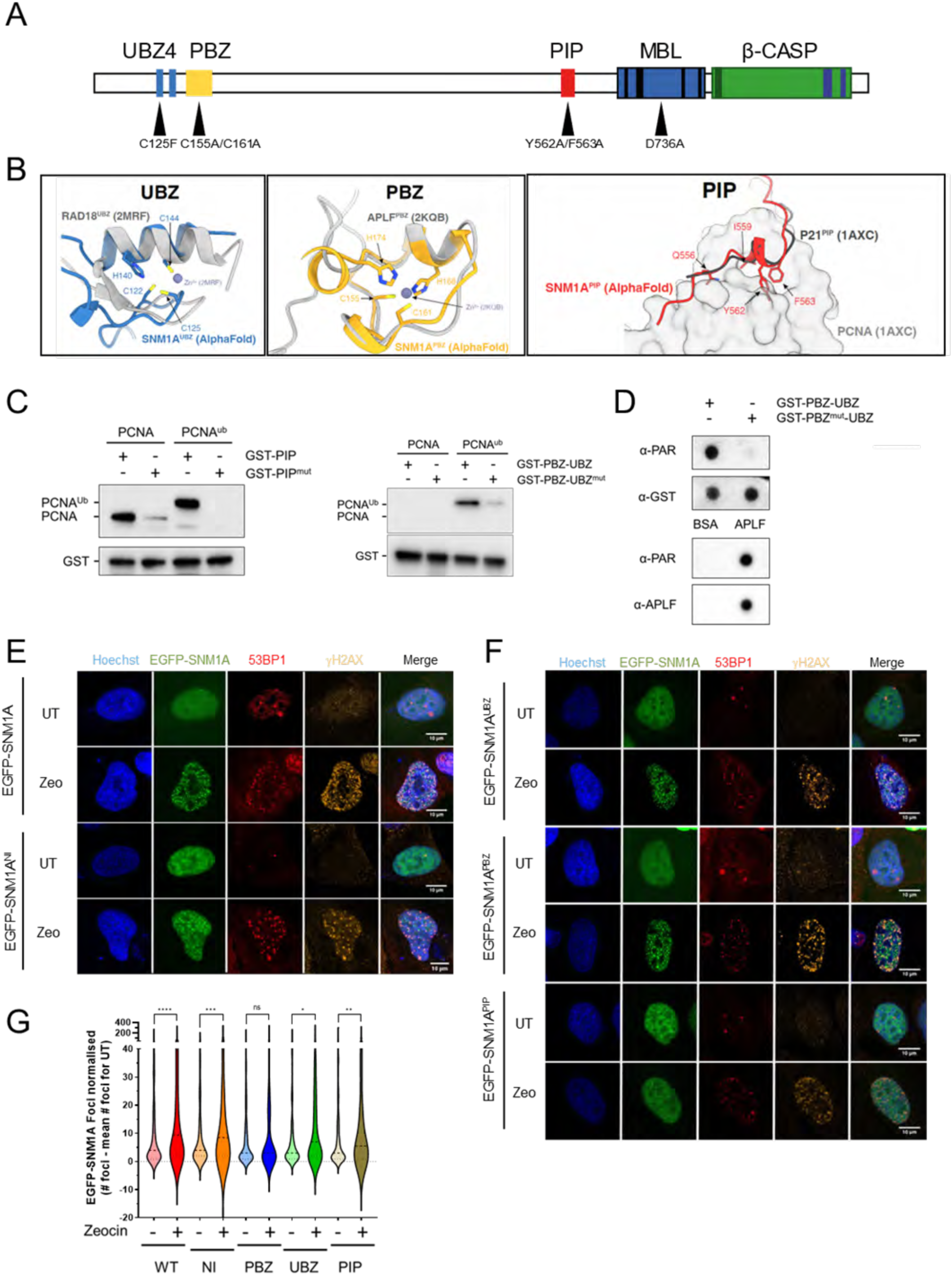
SNM1A contains conserved ubiquitin binding, PAR binding and PCNA interacting motifs. **A.** Schematic of domains and motifs of SNM1A, including location of residues predicted to disrupt ligand binding for each motif in the UBZ4 (ubiquitin-binding zinc-finger 4; C125F), PBZ (PAR-binding zinc-finger; C155A/C161A), PIP box (PCNA-interacting peptide; Y562A/F563A). **B.** Structural alignment of the UBZ (blue), PBZ (yellow) and PIP (red) domains using AlphaFold overlaid against their respective equivalents (in grey) from RAD18 (PDB 2MRF), APLF (PDB 2KQB) and P21 (PDB 1AXC) respectively. Highlighted amino acids reflect key conserved motifs which coordinate the Zn^2+^ for UBZ and PBZ, and fit into grooves within PCNA (1AXC) for the PIP box. **C.** GST-tagged SNM1A peptides were used to pull-down PCNA and lysine- 164 monoubiquitinated PCNA (PCNA^ub^). The wild-type PIP peptide (GST-PIP) binds and pulls down PCNA and PCNA^ub^, whereas the PIP box mutant (GST-PIP^mut^) does not (left-hand panel). A wild-type peptide spanning the region containing the PBZ and UBZ motifs (GST-PBZ-UBZ) pulls down PCNA^ub^, but not PCNA, and a peptide containing the mutated predicted UBZ binding residues (GST-PBZ-UBZ^mut^) fails to pull down PCNA and has a greatly reduced ability to pull down PCNA^ub^ (right hand panel). **D.** Slot blot of analysis SNM1A-GST peptides blotted to PVDF membrane indicate wild-type GST peptides spanning the PBZ and UBZ motifs (GST-PBZ-UBZ) are able to bind PAR chains (top panel), whereas those harbouring predicted interaction-disrupting mutations in the PBZ (GST-PBZ^mut^-UBZ) do not bind PAR chains. GST is probed as a loading control. The known PAR binding protein APLF1 was used as a positive control for PAR-chain binding whilst BSA is used as a negative control (bottom panels). **E.** Following Zeocin treatment SNM1A, 53BP1 and γH2AX foci form and co-localise at sites in both wild-type EGFP-SNM1A and also the nuclease inactive (NI) D736A mutant (EGFP-SNM1A^NI^). **F.** Protein harbouring substitutions of key residues in the UBZ, PBZ and PIP motifs (substitutions as shown in panel **A**.) show an altered recruitment of EGFP-SNM1A to Zeocin- induced sites of DNA damage. **G.** Quantification of these damage response EGFP-SNM1A foci (three biological repeats). Scale bars in E and F are 10 µm.

To explore these ligand interactions experimentally, we designed and produced a series of N-terminally GST-tagged peptides spanning the PBZ-plus-UBZ (residues 114 to 181) or PIP box (residues 547 to 575) motifs of SNM1A (Suppl. Fig. 5) and tested their capacity to interact with purified PCNA, purified lysine 164 ubiquitinated PCNA (PCNA^ub^) or PAR chains. The results revealed that wildtype PIP box peptides (GST-PIP) efficiently bind PCNA and PCNA^ub^, respectively, whereas a peptide containing substitutions at conserved residues in the relevant binding motifs, GST-PIP^mut^ (Y562A, F563A double substitution) did not pull down either PCNA or PCNA^ub^ (Fig. 3C, left-hand panel). While a GST peptide spanning the PBZ and UBZ motifs (GST-PBZ-UBZ) did not interact with native PCNA, PCNA^ub^ was pulled down by this peptide, indicating the UBZ motif-containing peptide can interact with ubiquitinated PCNA (Fig. 3C right-hand panel). A peptide harbouring structurally-predicted ligand binding mutation in the UBZ, a GST-PBZ-UBZ^mut^ peptide (C125F substitution), did not interact with PCNA^ub^ (or PCNA). This is consistent with the mutated UBZ residue mediating interaction with PCNA^ub^ as described above. Likewise, GST-PBZ-UBZ peptides were able to bind PAR chains in a dot-blot analysis, whereas GST-PBZ^mut^-UBZ (C155A and C161A double substitution) were not. Purified APLF, an established PBZ-containing protein^30^, acted as a positive control (Fig. 3D, lower panels).

Having confirmed that the PBZ, UPZ and PIP box mediate the predicted interactions *in vitro*, we investigated the potential role of each of these in the recruitment of SNM1A to complex DSBs. Cells expressing mutant forms of EGFP-SNM1A that ablate ligand interactions with PAR chains, ubiquitin and the PIP box, (EGFP-SNM1^PBZ^, double C155A, C161A substitution; SNM1A^UBZ^, C125F substitution; SNM1A^PIP^ Y562A, C161A double substitution, respectively) were expressed in U2OS cells and the induction of SNM1A and 53PB1 foci formation following Zeocin treatment was monitored (Fig. 3E and F) As before, wildtype EGFP-SNM1A protein colocalised with 53BP1 following Zeocin treatment; this localisation was independent of SNM1A catalysis, since cells expressing a SNM1A containing nuclease-inactivating mutations^17^ (harbouring a D736A substitution, here denoted SNM1A^NI^) was also efficiently recruited to the sites of Zeocin DSBs. Examination of the SNM1A^PBZ^, SNM1A^UBZ^ and SNM1A^PIP^ variants demonstrated a trend towards reduction in recruitment to Zeocin-induced foci that colocalised with 53BP1. The post-treatment increase in SNM1A foci was not statistically significant for the PBZ mutant, and exhibited reduced significance for the UBZ and PIP mutant forms, compared with the wildtype and nuclease inactive forms of SNM1A (Fig. 3G) which both exhibited significant post-treatment increase in foci.

Turning to a more directly quantifiable system, we used 405 nm laser microirradiation, without additional photosensitisers, as a sensitive and quantitative method to assess the relative contribution of the ligand-binding PBZ, UBZ and PIP motifs to the localisation and retention of SNM1A to complex DSBs. Direct 405 nm laser irradiation efficiently induces a high yield of single-and double-strand breaks associated with chemically modified termini, i.e. it acts as an effective surrogate for the damage induced by ionising radiation and radiomimetic drugs^37^. The EGFP-SNM1A foci in the damage tracks colocalised with 53BP1 foci produced, consistent with the efficient induction of DSBs by microirradiation (Fig. 4A). Examination of the kinetics of recruitment of EGFP-SNM1A in these cells revealed rapid accumulation of EGFP-SNM1A at damage sites, with apparent maximal accumulation within ∼10 minutes of irradiation (Fig. 4B and 4C). Having established a system to measure dynamics of EGFP-SNM1A at the sites of complex DSBs, we examined the impact of inactivation of the catalytic active site of SNM1A on recruitment dynamics. As with its ability to form foci following Zeocin damage, the EGFP-SNM1A^NI^ protein (SNM1A^D736A^) was recruited to laser stripes with kinetics indistinguishable to the wildtype protein, confirming that the catalytic activity of SNM1A can be separated from its recruitment to and retention at complex DNA breaks (Fig. 4B and 4C). Next, we investigated the possibility of cell cycle phase-dependency on recruitment of EGFP-SNM1A to complex DSBs. EGFP-SNM1A was rapidly recruited to laser induced damage in G1-, S- and G2-phase cells (Fig. 4D, E), implying a role for SNM1A in processing complex DNA breaks that is sustained throughout the cell cycle.

**Figure 4.**
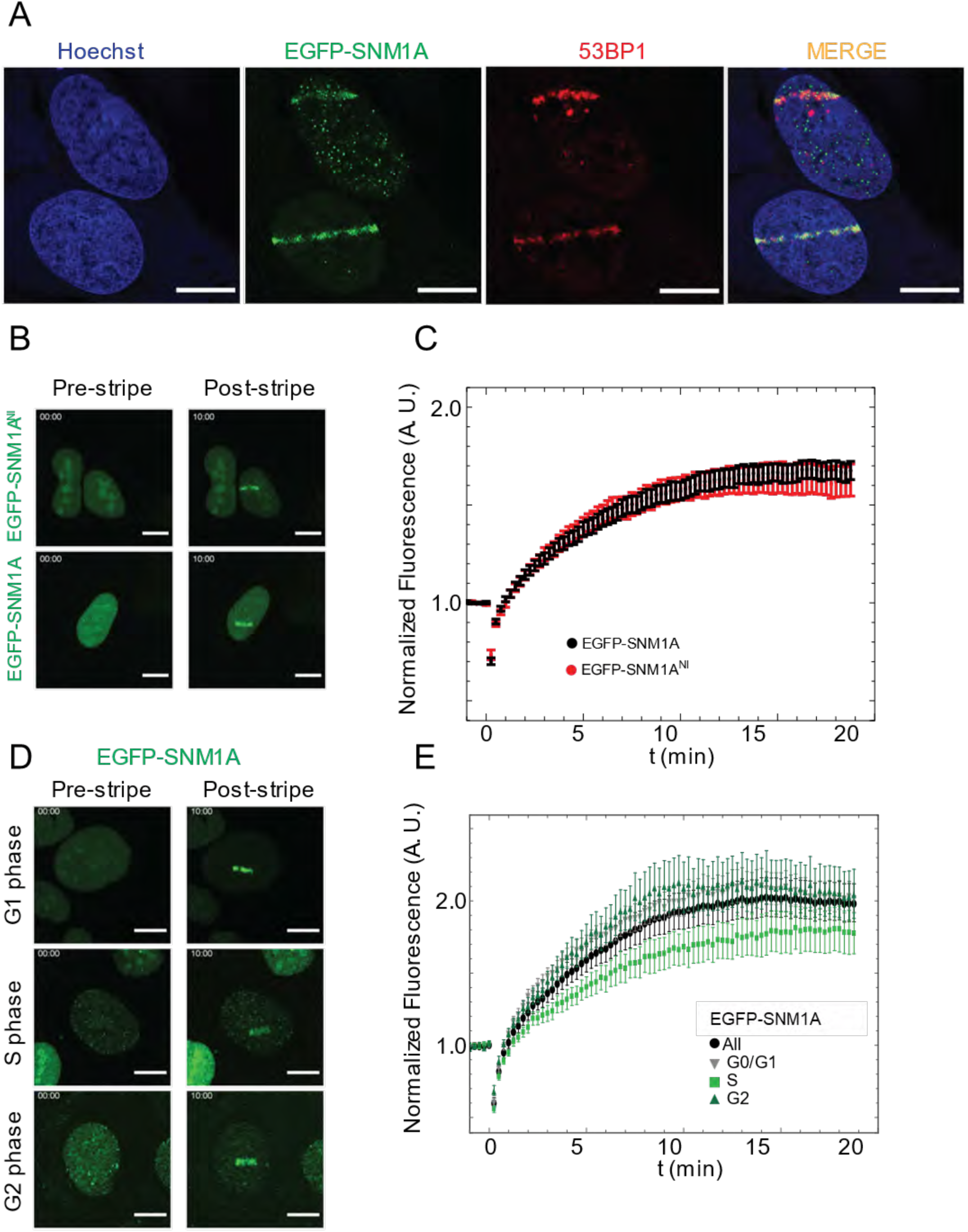
SNM1A is rapidly recruited to laser-induced DNA damage. **A.** EGFP-SNM1A is recruited to localised laser induced DNA damage (laser stripes) where it colocalises with 53BP1. Staining with Hoechst 33258 was used to define nuclei. **B.** To analyse the recruitment of EGFP-SNM1A cells were monitored over 20 minutes and the fluorescence intensity at the sites of laser striping measured (see Material and Methods). Both wild-type EGFP-SNM1A (wild-type) and nuclease inactive EGFP-SNM1A^NI^ were recruited to laser damage with similar kinetics and intensity (quantified in panel **C**). **D.** Recruitment of EGFP-SNM1A was similar in G0/G1, S and G2-phases of the cell cycle (quantified in panel **E.**). All laser striping experiments represent data from at least ten cells from three biological repeats, error bars are standard error of the mean. Scale bars in A, B, and D are 10 µm.

### The PBZ, UBZ and PIP box domains collectively mediate recruitment and retention of SNM1A at complex breaks

We then analysed the roles of the UBZ, PBZ and PIP box motifs in the recruitment of SNM1A to complex DSBs (Fig. 5A and 5B). For both EGFP-SNM1A^PBZ^ and EGFP-SNM1A^PIP^, delayed recruitment was observed and these proteins never achieved the same local concentration at the laser site observed for the wildtype SNM1A. By contrast, the EGFP-SNM1A^UBZ^ exhibited a near-normal initial rate of recruitment, though not reaching the levels observed for wildtype SNM1A, followed by gradual loss from the laser sites. EGFP-SNM1A^PIP^ exhibited the most dramatic initial defect in laser stripe recruitment, suggesting that PCNA interaction is particularly important for SNM1A recruitment to complex breaks (Fig. 5A, B; representative movies are shown in Suppl. Fig. 6A). Overall, these observations suggest a key role for the PIP box in the initial recruitment of SNM1A to laser damage sites, with the PBZ also making a contribution to initial recruitment, and that the UBZ motif acts to stabilise the recruited protein at complex DSBs.

**Figure 5.**
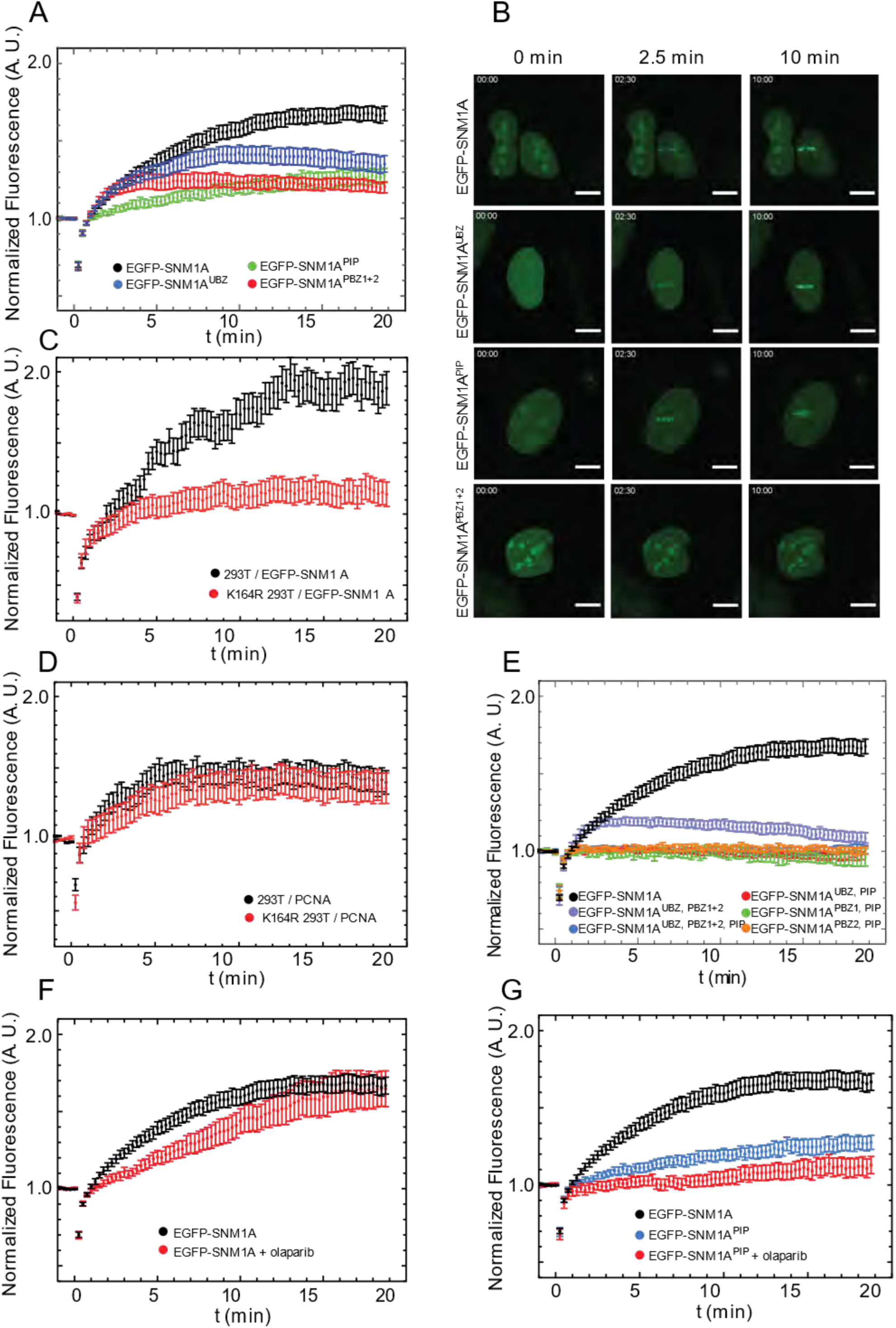
The UBZ, PBZ and PIP box motifs of SNM1A play a role in protein recruitment to sites of laser- induced DNA damage. **A.** The recruitment of EGFP-SNM1A containing substitutions in the UBZ (C125F), PBZ (C155A and C161A) and PIP box (Y562A and F563A) that ablate direct ligand-binding (see Figure 3) were monitored following laser-induced damage, representative images shown in panel **B. C.** Transiently expressed EGFP-SNM1A recruitment to laser stripes in HEK293T and K164R 293T cells (cells expressing a mutation in the ubiquitin modified PCNA residue which are unable to be ubiquitinated on K164). **D.** Recruitment of PCNA to laser stripes as measured through a transiently expressed RFP-PCNA chromobody in HEK293T and K164R HEK293T cells. **E.** Double or triple co-substitution of key residues in the UBZ, PBZ and PIP box were used to investigate the cooperative contribution of these SNM1A motifs to laser stripe recruitment. **F**. Olaparib treatment reduces EGFP-SNM1A recruitment to laser stripes. **G.** The effect of combining PIP box mutation with Olaparib treatment on laser stripe recruitment. All laser striping experiments represent data from at least eight cells from a minimum of three biological repeats, error bars are standard error of the mean. Scale bars in B are 10 µm.

We also investigated the kinetics of PCNA recruitment to complex DSBs produced by the 405 nm laser, creating a cell line that stably expressed both EGFP-SNM1A and an anti-PCNA RFP-tagged nanobody^38^ (Chromobody®, anti-PCNA VHH fused to red fluorescent protein). We observed that PCNA is recruited to the sites of Zeocin-induced foci and to the sites of laser stripes, where it colocalises with EGFP-SNM1A (Suppl. Fig. 7A, B, C with representative movie shown in Suppl. Fig. 6B). PCNA recruitment to DSBs precedes that of EGFP-SNM1A by several minutes (Suppl. Fig. 7B, C), occurs in any phase of the cell cycle (Suppl. Fig. 7D) and PCNA recruitment is not delayed or reduced in SNM1A^-^ cells (Suppl. Fig. 7E).These observations imply a key role for PCNA in attracting EGFP-SNM1A to complex DSBs.

To further examine the role of the UBZ, we examined the recruitment kinetics of EGFP-SNM1A in cells depleted for the key E3 ubiquitin ligases involved in DSB repair. These include RAD18 (which is an established E3 ligase for the ubiquitination of PCNA)^39^, RNF8 and RNF168 which mono- and poly-ubiquitinate histone H2A, respectively, in response to DSBs. Depletion of these three E3 ligases using siRNA revealed that only depletion of RAD18 impacted SNM1A recruitment to laser damage, and the defect was kinetically similar to mutation of the UBZ domain since it led to wild-type-like initial recruitment kinetics of SNM1A to the stripes, but lower overall levels of accumulation at these sites (Suppl. Fig 7F). The only known target of RAD18 E3 ligase activity is lysine 164 of PCNA, and indeed a wild-type UBZ motif directly interacts with PCNA^ub^ *in vitro* (Fig. 3C). However, we were unable to detect monoubiquitination of lysine 164 by immunoblotting in whole cell extracts, using a ubiquitin-specific antibody raised against this epitope, likely because the fraction of PCNA which is ubiquitinated in these cells is low. A functional ubiquitin response is clearly required for the efficient recruitment and retention of SNM1A at the sites of laser damage, since pre-treatment of cells with MG132, an agent that exhausts the cellular free ubiquitin pool by proteasome inhibition^40^, dramatically reduced recruitment of EGFP-SNM1A to laser stripes (Suppl. Fig. 7G). Nonetheless, to definitively address this point, we employed edited 293T cells where lysine 164 has been substituted with an arginine residue (K164R)^41^. Here, EGFP-SNM1A recruitment to laser stripes was delayed in a manner the phenocopies mutation of the UBZ motif (compare Fig. 5C to Fig. 5A), while PCNA recruitment was unaffected by mutation of this residue, as determined by Chromobody® detection (Fig. 5D).

To examine the interplay and interdependence of the three key conserved motifs involved in SNM1A break localisation, we created double-mutations in the PBZ and UBZ, PBZ and PIP box, and UBZ and PIP box motifs, and a form of SNM1A triply mutated in the PBZ, UBZ and PIP box. Analysis of the dynamics of recruitment and retention of the double- and triple-mutated forms of SNM1A reveals that mutation of either the PBZ or UBZ motifs together with the PIP box essentially eliminated SNM1A recruitment to and retention at laser stripes, where dual mutation of the PBZ and UBZ drastically reduced recruitment and retention of EGFP-SNM1A to stripes (Fig. 5E). Accordingly, mutation of all three motifs eliminates recruitment (Fig. 5E).

To explore the role of the PBZ in recruitment of SNM1A to complex DSBs, we exploited, Olaparib which competitively inhibits PARP1 and also produces PAR chain-shielding in cells through trapping the PARP enzyme during catalysis^42^. Pre-treatment of cells with Olaparib led to a reduced rate of recruitment of EGFP-SNM1A to sites of laser damage (Fig. 5F). Combining Olaparib treatment with the EGFP-SNM1A^PIP^ mutant led to an abrogation of recruitment and retention reminiscent of the double substitution mutations in the PBZ and PIP motifs (Fig. 5G). This observation implies that interaction with PAR chains is important for efficient initial recruitment of SNM1A to laser damage. The combined results demonstrate that several conserved motifs work collectively to orchestrate the initial recruitment (PBZ and PIP) of SNM1A to complex breaks and that the UBZ motif is important for SNM1A retention at these breaks, likely in a manner involving association with PCNA^ub^.

Since multiple motifs in SNM1A act in concert to recruit and retain SNM1A at sites of complex DNA damage, we investigated whether interactions with their cognate ligands impact on the exonuclease activity of SNM1A. Although the core catalytic domain of SNM1A is formed by the C-terminally located MBL-β-CASP fold, we wanted to investigate whether the N-terminal motifs that regulate the damage localisation of SNM1A (PBZ, UBZ and PIP) and of SNM1A impact on catalytic activity. To do this, we used a kinetically preferred substrate of SNM1A^43^, a single-stranded 21-nucleotide (21-nt) oligonucleotide, bearing a 5ʹ-phosphate group and radiolabelled at its 3ʹ-end. When incubated with full-length SNM1A (which was purified from human cells harbouring the ligand-binding motif substitutions utilised in the preceding cellular studies) all forms of the protein retained nuclease activity, with the caveat that purification of full-length SNM1A yields small quantities of protein and some variation in activity from preparation to preparation was observed (Suppl. Fig. 8A).

We next tested the impact of PCNA and PAR chains on the activity of full-length SNM1A. Titration of PCNA followed by analysis of the digestion activity over a time course of one hour revealed that PCNA did not enhance the activity of SNM1A, with some decrease in digestion being observed (Suppl. Fig. 8B). To determine whether this was a direct effect mediated by interaction of PCNA and SNM1A, we repeated this experiment with N-terminally truncated SNM1A (ΔN-SNM1A, residues 697-1040) retaining the MBL-β-CASP fold (and exonuclease activity), but lacking the PIP box. ΔN-SNM1A activity was also reduced in the presence of PCNA (Suppl. Fig. 8C) suggesting that perturbation of nuclease activity in the presence of PCNA is likely a result of reduced SNM1A access to the 5ʹ-terminus of its DNA substrate in the presence of PCNA. Next, we incubated SNM1A with PAR chains, or performed reactions in a system that produces PAR chains *in situ* by pre-incubating poly-ADP-ribose polymerase with NAD and the DNA substrate with subsequent addition of purified full-length SNM1A to the reaction. The activity of SNM1A was not modulated by the presence of PAR chains (Suppl. Fig. 8D). Together, these biochemical analyses suggests that the interactions mediated by the PBZ, UBZ and PIP box motifs are principally important for recruiting and retaining SNM1A to DNA damage, rather than modulating the activity of the enzyme.

### SNM1A can process DNA containing oxidised lesions

It was of interest to investigate whether the requirement of SNM1A for efficient repair of complex DSBs relates to the ability of SNM1A to process DNA containing lesions induced by radiation and radiomimetic damage, as proposed for its established role in resecting DNA containing ICLs. We initially employed 21-mer double-stranded oligonucleotides that contain an 8-oxoguanine (8-oxo-G) residue on the substrate strand, one of the major oxidative lesions induced by radiation at complex breaks^44^, either located centrally, or at the 5ʹ-terminus. Incubation of these substrates with ΔN-SNM1A revealed that its exonuclease activity can traverse the lesion without pause or arrest (Fig. 6A). This striking activity contrasts starkly with that of human Exonuclease 1 (EXO1), one of the major exonucleases involved in performing DNA extensive end resection in preparation for homologous recombination, which is quantitatively arrested by an 8-oxo-G lesion, regardless of whether it is located terminally or centrally in the substrate (Fig. 6A; baseline activities of SNM1A and EXO1 were normalised on an undamaged substrate, Fig. 6A, first 3 lanes). We examined the capacity of SNM1A to process additional oxidative lesions commonly associated with complex DSBs. Examining related substrates that contain thymine glycol and hypoxanthine bases reveals that SNM1A efficiently digest ssDNA substrates containing these lesions, with kinetics comparable to those observed on a native, undamaged template (Fig. 6B), and again, we observed thymine glycol acted as a complete block to EXO1 digestion (Suppl. Fig. 8E). Finally, we determined whether SNM1A can process DNA substrates containing Zeocin-induced DNA breaks *in vitro*. We employed two doses of Zeocin in an iron-metal catalysed reaction that mimics the activation of Zeocin in cellular conditions^45^, we introduced SSBs (0.05 mg/mL Zeocin; producing relaxed open-circular DNA) and a mixture of SSBs and DSBs (0.5 mg/mL Zeocin; producing relaxed open-circular DNA and linear DNA forms) into plasmid DNA. SNM1A was capable of extensively digesting all of the nicked and linear forms of DNA molecule (Fig. 6C), consistent with a capacity to exonucleolytically process the chemical modifications induced by radiomimetics at the sites of the DNA breaks they induce.

**Figure 6.**
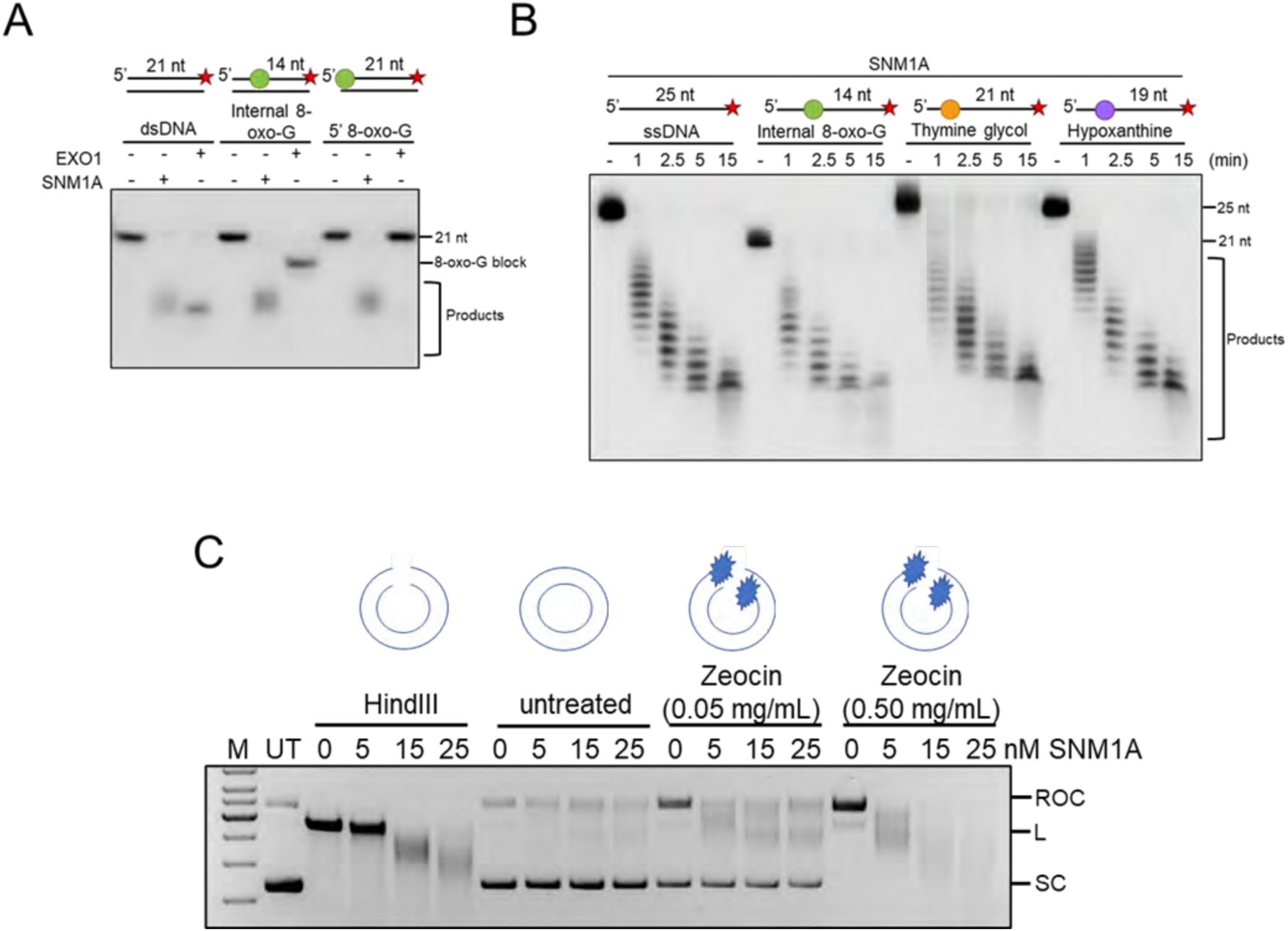
Purified SNM1A can digest complex DNA lesions in vitro. **A.** Purified SNM1A and Exo1, can digest 3′ radiolabelled dsDNA *in vitro.* SNM1A and Exo1 were also incubated with oligonucleotides with either an internal or a 5′-located 8-oxo-guanine (8-oxo-G). SNM1A can digest past the altered 8-oxo-G bases but Exo1 does not. **B.** SNM1A can digest substrates containing several major oxidative DNA damage products, including 8-oxo-G, Thymine glycol and Hypoxanthine. **C.** Untreated (UT) or HindIII linearised pG46 plasmid was treated in the presence or absence of Zeocin (0.05 mg/mL and 0.5 mg/mL) in a pharmacologically-relevant iron-catalysed reaction to induced complex DNA breaks. Plasmids were treated with purified SNM1A (0 to 25 nM) and the resulting DNA products resolved on an agarose gel, stained with Ethidium bromide and visualised. SC = supercoiled DNA; ROC = Relaxed Open-Circular DNA; L = Linear DNA.

## Discussion

Disruption of SNM1A in human cells reveals an unanticipated phenotype, a profound sensitivity to radiomimetic drugs and IR treatment, that is associated with delayed resolution of DSBs. SNM1A is recruited to the complex DNA breaks induced by radiomimetics, IR and long-wave laser damage within minutes of damage induction. Multiple potential ligand-interacting motifs are present in the N-terminal region of SNM1A, proximal to the MBL-β- CASP domain which forms the catalytic core and active site^15^. Systematic examination of the role of the SNM1A putative interaction and recruitment motifs revealed an important role for the PIP box and PBZ motif in efficient initial recruitment of SNM1A to sites of complex breaks. This finding implies that interaction of SNM1A with PCNA (which is itself very rapidly recruited to laser-induced damage, as demonstrated here and by others^46^) and PAR chains is critical for timely recruitment of SNM1A to laser damage. This conclusion is consistent with the reported ICL-induced interaction of SNM1A with PCNA^18^; the new data presented here demonstrates that SNM1A interacts with PAR-chains *in vitro* and in a PBZ-dependent manner. Moreover, trapping and inhibiting poly-ADP-ribose polymerase (PARP) with the PARP inhibitor Olaparib phenocopies mutation of the PBZ motif, i.e. a reduction in initial SNM1A recruitment to laser damage is observed. Indeed, co-disruption of the SNM1A PIP box with the PBZ completely abrogates recruitment of SNM1A to the sites of laser and Zeocin-induced breaks and, consistently, so does mutation of the PIP box when combined with Olaparib treatment.

SNM1A contains a UBZ motif, adjacent to its PBZ motif. Like its PIP box motif, the UBZ motif is reported to be important for targeting SNM1A to the ICL damage foci during S-phase^18^. In response to laser-induced damage, mutation of the UBZ motif produced a phenotype distinct from that observed by mutation of the PBZ or PIP box. Importantly, this result suggests that mutations in the neighbouring UBZ or PBZ motifs do not mutually impact on the function of the other motif. UBZ mutant SNM1A is recruited with near-normal initial kinetics to laser damage, but fails to accumulate to the same final level. A screen of ubiquitin E3 ligases that are known to deposit ubiquitin at sites of DNA breaks, and which are therefore candidates for providing the ligand for the interaction with the SNM1A UBZ motif, reveals that RAD18 loss phenocopies mutation of the UBZ, leading to reduced overall levels of accumulation of SNM1A at laser stripes. PCNA (lysine 164) remains the only known target for RAD18 ubiquitination^47^, though we were unable to detect PCNA ubiquitination following IR or Zeocin treatment, either in whole cell extracts or by performing immunoprecipitations with an antibody directed against monoubiquitinated PCNA. This likely because the fraction of PCNA that is ubiquitinated is low. A role for RAD18 in DSB and replication fork repair that relies on interaction with the SLF1 and SLF2, and is important for recruitment of the SMC5/6 complex to chromatin, has been previously reported^48^. However, this RAD18-subcomplex is recruited to damage by RNF168, which is dispensable for SNM1A recruitment to DSBs (Suppl. Fig. 7), suggesting that the RAD18-SLF1-SLF2 complex does not play a major role in recruiting SNM1A to DSBs. Moreover, as cellular PCNA K164 is required for normal SNM1A retention at lasers stripes (Fig. 5C), and the UBZ motif of SNM1A directly interacts with ubiquitinated PCNA^ub^ (Fig. 3C), the evidence that ubiquitinated PCNA acts to retain SNM1A at the sites of such damage is extremely robust.

The phenotypes of cells disrupted for homologues of SNM1A has been examined in multiple organisms, ranging from yeasts to humans, and has revealed a conserved role in the repair of ICLs. However, where the sensitivity and a response to IR and radiomimetics has been examined in vertebrate cells, only human cells have been implicated in response to these forms of damage. In the case of mouse ES cells and chicken DT-40 cells no marked sensitisation to IR was observed^49–51^. However, work from Richie and colleagues has shown that human SNM1A forms subnuclear foci following IR treatment^52^, and that these foci co-localise with 53BP1 and Mre11, providing early evidence for a role for SNM1A in repair of radiation-induced DSBs. A potential explanation for these interspecies differences rest with the fact that vertebrates harbour (at least) three SNM1 paralogues, SNM1A, SNM1B/Apollo and SNM1C/Artemis. Based upon substantial similarities in their 5ʹ-3ʹ exonuclease catalytic activities, SNM1B might plausibly play a related or redundant role in the processing of radiation-induced DSBs. Indeed, multiple reports indicate that loss of murine SNM1B is associated with increased radiosensitivity^20, 53^. Therefore, in mice the roles of SNM1A and SNM1B in the repair of complex DNA breaks might only be revealed once such redundancy or species-specific prioritisation of their roles has been systematically examined. Moreover, Artemis/SNM1C deficient cells are IR sensitive and Artemis/SNM1C plays an established role in removal of several end-blocking chemical modifications during repair of complex breaks. Strikingly, disruption of the SNM1A homologue in budding yeast, Pso2, together with inactivation of the nuclease Mre11 (*via* an *mre11-H125N* mutation) leads to a profound sensitivity to IR, suggesting that these two nucleases may play, at least partially, redundant roles in processing complex DNA ends^54^. Notably, like the *pso2* and *mre11* single mutants, *pso2 mre11-H125N* double mutant cells did not display any overt defects in the repair of ʹclean DSBsʹ induced by the HO-endonuclease, analogous to the situation we report here for SNM1A in the I-SceI reporter assay (Suppl. Fig. 4). This suggests that the key function of Pso2 and SNM1A (and a known major function of Mre11) is in processing of chemical modifications to DNA at ʹdirtyʹ break termini prior to their repair, rather than direct participation in the DSB process *per se*. MRE11 is also implicated in the initiation of DSB resection and repair cells by clearing covalently linked proteins at break termini, for example the DNA-topoisomerase 1 (Top1) crosslinks induced through camptothecin (CPT) treatment^55, 56^. However, we see no evidence of a role for SNM1A in this process, based on the wildtype like sensitivity of SNM1A cells to CPT. It is also interesting to note that co-inactivation of Pso2 with Exo1, the only other known major 5ʹ-3ʹ repair exonuclease in yeast, does not strongly impact on IR sensitivity (our unpublished observations and^54^). This observation implies that Exo1 is not a major factor acting to process chemically modified termini in the absence of Pso2, and *vice versa*. Our finding that the exonuclease activity of human EXO1, a close functional relative of yeast Exo1, is highly sensitive to the presence of chemical lesions is consistent with this proposal. It seems likely that both MRE11 (through its 3ʹ-5ʹ exonuclease or endonuclease activities) and SNM1A (through its 5ʹ-3ʹ exonuclease activity) are well equipped to deal with a set of the chemical lesions induced at break termini by radiation and play a major role in this important defence against complex DSBs. Therefore, a comprehensive examination of the relationship between all three SNM1 paralogues in mammalian cells, in addition to their contribution relative to other established and putative end-processing nucleases (in particular MRN), is warranted in future studies.

Finally, the identification of a role for SNM1A in processing radiation-induced oxidative modified termini in DSBs raises the possibility that SNM1A inhibition might be used to reduce the doses of radiation treatment required to treat cancer. The results presented here show that the interactions mediated by the PBZ, UBZ and PIP box motifs are principally and collectively important for recruiting and retaining SNM1A to DNA damage, rather than modulating the activity of SNM1A. Thus, targeting the β-CASP-MBL fold domain of SNM1A is likely a preferred mode of inhibition and is the subject of ongoing efforts.

## Acknowledgements

This work was supported by a Cancer Research UK Programme Award (CRUK/ A24759) to PJM and CJS. MR is in receipt of an MRC Graduate Studentship, LRH has received a Wellcome Trust Studentship and SL was supported by a National Science Scholarship from the Singaporean Agency for Science, Technology, and Research (A*STAR). We thank Opher Gileadi for purified APLF, Paul Modrich for EXO1 constructs, Philip Hublitz for help with CRISPR guide design, and Andrew Blackford for sharing XRCC4 disrupted cells. George-Lucian Moldovan kindly provided 293T PCNA^K164R^ cells.

## Contributions

LPS, BCL, and DW performed cell biology experiments. HTB, LRH, BS, SL, ACM and JN undertook protein purification and biochemical assays. MR produced the AlphaFold models and figures and undertook data analysis. CR and HDU provided PCNA and PCNA^ub^ reagents and expertise. LPS conduced all other experiments. CJS and PJM supervised the project. LPS and PJM wrote the manuscript.

## Materials and Methods

### Cell lines

U2OS (American Type Culture collection: HTB-96), 293FT (ThermoFisher Scientific: R70007), HEK293 XRCC4^-^ (a gift from Andrew Blackford) and 293T PCNA^K164R^ cells (a gift of George-Lucian Moldovan) cells were cultured in D-MEM medium supplemented with 10% foetal bovine serum.

### Creation of SNM1A^-^ cell lines using genome engineering

Zinc finger nucleases (ZFN) and CRISPR-Cas9 genome editing technologies were employed to make stable deletion-disruptions in *DCLRE1A*, the gene encoding SNM1A. CompoZr ZFNs were designed by and purchased through Sigma-Aldrich with a detection and cut site in the *DCLRE1A* gene as follows (cut site in lowercase): TGCCAGATGCCTTTTtcctcATTGATAGGGCAGAC

Cells were transfected with plasmids (∼1.3 µg) containing forward and reverse ZFNs in 10 µL of Lipofectamine 2000 in a total volume of 200 µL OptiMEM media (Sigma). The transfection mixture was incubated (10 min) to allow the complexes to form before being added to 5 x 10^5^ cells that were freshly plated in 2 mL of medium in wells of a 6-well plate. The medium changed after 12-18 hours; after a further 72 hours, the cells were analysed for genomic insertions or deletions using the SURVEYOR Mutation Detection Assay (Integrated DNA Technologies, Cel I assay). Once a pooled cell population was identified to contain cells with altered genomes, dilution cloning was performed. Once grown, the clones were assayed for alterations to their genomes through genomic PCR. Identified SNM1A^-^ clones were validated by analysing SNM1A protein levels (through SDS-PAGE/Western blot techniques) and mRNA levels (through defined qRT-PCR).

CRISPR-Cas9 genome editing was performed using sgRNAs designed using an in-house design tool. The targets for the CRISPR-Cas9 sgRNAs of the *DCLRE1A* gene were as follows (the cuts remove the majority of exon 1 of the *DCLRE1A* gene):

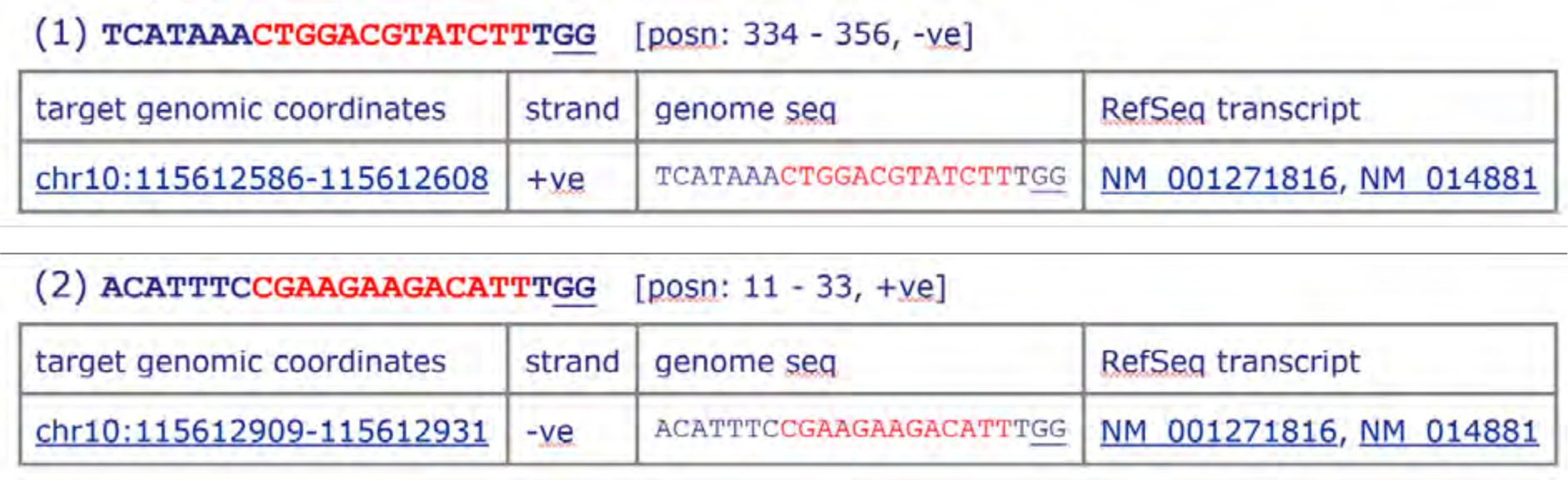

The sgRNAs were cloned in to CRISPR-Cas9 vectors pX330 and pX458 carrying GFP and RFP fluorescence expression markers respectively.

Once treated with both plasmids (as above for ZFNs), the cells were sorted by FACS to collect individual GFP and RFP positive cells in a well of a 96 well plate. The clones were grown before being analysed for genomic alterations (as above). The SNM1A^-^ cells used are detailed in Suppl. Fig. 1.

### Cell transfections for expression

For laser microirradiation, or analysis of drug induced DNA damage (foci), cells were transfected with pEGFP-C1 plasmids containing wildtype SNM1A and substitutions in the domains of interest. Cells (5 x 10^5^) were seeded in 35 mm glass bottom “μ-Dish” dishes (Ibidi). Transfection was performed with ∼2.6 µg plasmid in 2 mL media with 10 μL Lipofectamine 2000 (ThermoFisher scientific). Complexes were allowed to form for 10 minutes and then added dropwise to the cells.

For creation of cells stably expressing plasmids containing EGFP-SNM1A or a Chromobody^®^ of RFP-PCNA (Chromotek), ∼5 x 10^5^ cells were seeded in wells of a 6-well plate and immediately transfected with the desired plasmid (2.6 µg) with Lipofectamine 2000 (2 µL) in a total volume of 200 µL. A reduced volume of Lipofectamine 2000 was used to reduce the toxicity during transfection. After 3 h, the media was changed and the cells were grown for 24 hours. The cells were sorted and single cells expressing the fluorescence marker associated with the plasmid were plated out in a 96 well plate and allowed to grow until 50-75% confluent. The cells were transferred to 25 cm^2^ flasks and allowed to grow further. Once 60-80% confluent, cells were analysed for fluorescence and clones that exhibited adequate level of fluorescence were further analysed by microscopy, SDS-PAGE, and western blot protocols to establish the desired expression and expected phenotype of the clones. For PIP box, PBZ and UBZ substitution mutants, standard site-directed mutagenesis techniques were used to introduce the indicated sequence changes.

Cells that had successfully integrated the pEGFP-C1-SNM1A plasmid into their genome were first selected with kanamycin before being sorted by FACS into single wells in a 96 well plate, and grown up as a clonal population.

### BRCA2 siRNA Transfections

Cells were treated with 20 nM siRNA against BRCA2 (GGGAAACACUCAGAUUAAAUUdTdT and AAUUUAAUCUGAGUGUUUCCCdTdT, Life Technologies) in HiPerFect transfection reagent (Qiagen) using a fast-forward transfection protocol then further siRNA is added 24 hours later, as per the manufacturer’s protocol.

### Cell treatments with DNA damaging drugs and radiation

Cell treatments, unless otherwise stated in figure legends, were; Cisplatin (CDDP, Teva pharmaceutical industries Ltd., Eastbourne, UK. Cat # 51642169) is a concentrate for clinical infusion, including 1mg/mL cisplatin with sodium chloride, hydrochloric acid/sodium hydroxide (for pH adjustment). If required, CDDP was diluted in PBS, treatments were for 4h. Zeocin (Invitrogen, Thermo Fisher Scientific, Cat # R25001) was diluted in PBS. Treatments were at 0.1mg/mL for 2h, unless otherwise listed in figure legends. Ionising radiation induced damage (IR) was induced using a caesium 137 source. SJG-136 (a kind gift from John Hartley, UCL) was dissolved in H_2_O and treatments were performed continuously. Olaparib was made up in PBS and clonogenic treatments were continuous. Methanol free aqueous formaldehyde (16% (v/v) (Taab Laboratories Equipment) was always freshly diluted in PBS and used immediately. UVC treatments were performed using a Stratagene, UV stratalinker 2400 machine with UV-C lamps. Prior to UV treatment, the media was removed from the dishes, the cells were dosed with UVC and fresh media added. Hydrogen peroxide was diluted in PBS from a stock 30% solution (BDH, Cat#BDH7741-1). MG-132 (Med Chem Express) was dissolved in DMSO and cells were treated with 5 μM. Camptothecin (Cambridge Bioscience) was dissolved in DMSO.

### Colony counting/clonogenic assays

Clonogenic assays were performed in 10 cm tissue culture dishes (for U2OS and clones) or T75cm^2^ tissue culture flasks (293FT cells, due to their reduced colony forming ability). Cells (1000 per dish for U2OS cells and 2000 per flask for 293FT cells) were seeded in complete media (10 mL and 20 mL respectively) and allowed to attach overnight before being treated with the desired agents. Following treatments (short or continuous) cells were allowed to grow and form colonies for 10 days. Colonies formed were stained with Coomassie R250 (Sigma) and counted on a COLCount Colony Counter (Oxford Optronix). All experiments represent the mean (±SEM) of at least three biological repeats of duplicate dishes/flasks for each treatment.

### Structural prediction

ColabFold Google Colabs notebooks were used to predict structures of the UBZ, PBZ and PIP Box domains (https://colab.research.google.com/github/sokrypton/ColabFold/blob/main/AlphaFold2.ipynb)^57^. With default parameters and inputting the sequences: (numbering as for human SNM1A; NCBI /NP_001258745.1/): UBZ = 112-148; PBZ =153-182; and PIP = 552-573. Coordinates of the highest-ranking AlphaFold models were then visualised, aligned to prior structures, and rendered using ChimeraX^58^. Alignments were made to prior structures as follows: RAD18 UBZ – PDB 2MRF^34^; APLF PBZ – PDB 2KQB^35^; PCNA with P21 PIP – PDB 1AXC^36^.

### DNA content and cell cycle FACS analyses

An estimation of distribution of cell cycle phase of treated cells was performed using BrdU incorporation and DNA content analysis as described^17^. Quantification of the phase of the cell cycle was performed post acquisition using FCS Express software (De Novo Software), example gates used are shown in Suppl. Fig. 3A.

### Laser damage striping

DNA damage microscopy experiments were conducted using a Zeiss 780 or Zeiss 880 inverted confocal microscopes using a Plan-APO 63X 1.40NA oil immersion objective, with the optical zoom adjusted to a projected pixel size of 100 nm, at physiological conditions (37°C and 5% CO2). Prior to all experiments, care was taken to adjust the collimation of the 405 nm laser such that the z-focus aligned with the visible lasers (488, 561, and 633 nm) as determined by imaging of 200 nm diameter TetraSpeck beads. The laser power of the 405 nm lasers on both the Zeiss 780 and the Zeiss 880, as measured at the focal plane of a 10X 0.45NA air objective, was about 5 mW.

DNA damage was induced by defining a rectangular ROI damage site within the cell nuclei with varied length (6.5 μm for single cell experiments and around 200 μm for multi-cell experiments) but fixed width of 1 μm. The laser damage was then inflicted using the 405 nm at full laser power, a scan speed pixel dwell time setting of 8 μs, and with 25-line iterations. To observe recruitment of EGFP tagged wildtype and mutant SNM1A to the DNA damage sites, we performed time-lapse experiments for a total of 80 image frames at a frame interval of 15 sec where the 405 nm laser induced DNA damage was initiated after image frame 5. EGFP emission in these experiments were collected on GaAsP detector, with a pinhole setting of 1 AU, a bandpass emission setting of 499-544 nm, a pixel dwell time of 1.58 μs, and with a line averaging setting of 4.

Quantitative analysis of the active recruitment to the DNA damage site was performed using Image J and Mathematica. We extracted the mean fluorescence intensity F(t)Bleach from a region-of-interest (ROI), superimposed on the DNA damage site, relative to the mean fluorescence intensity at a site on the same cell but away from the damage site F(t)Control) as a function of time, from several cells. The data were normalized by the fluorescence intensity F(-) prior to the induction of the DNA damage such that active recruitment results in a ratio of [F(t)/F(-)]Bleach / [F(t)/F(-)]Control > 1 versus no recruitment or passive recruitment results in a ratio of [F(t)/F(-)]Bleach / [F(t)/F(-)]Control <= 1. The presented normalized data represents the mean (+/- SEM) for at least n=10 cells from at least three biological repeats.

Cells were scanned in a variety of phases of the cell cycle. We determined the cell cycle phase using multiple approaches. PCNA forms foci during DNA replication, which indicates the cells are in S-Phase. Cell morphology coupled to determining mean DAPI volume was used to identify G1 and G2/M cell phases, respectively, as described^59^.

### List of antibodies used in this study

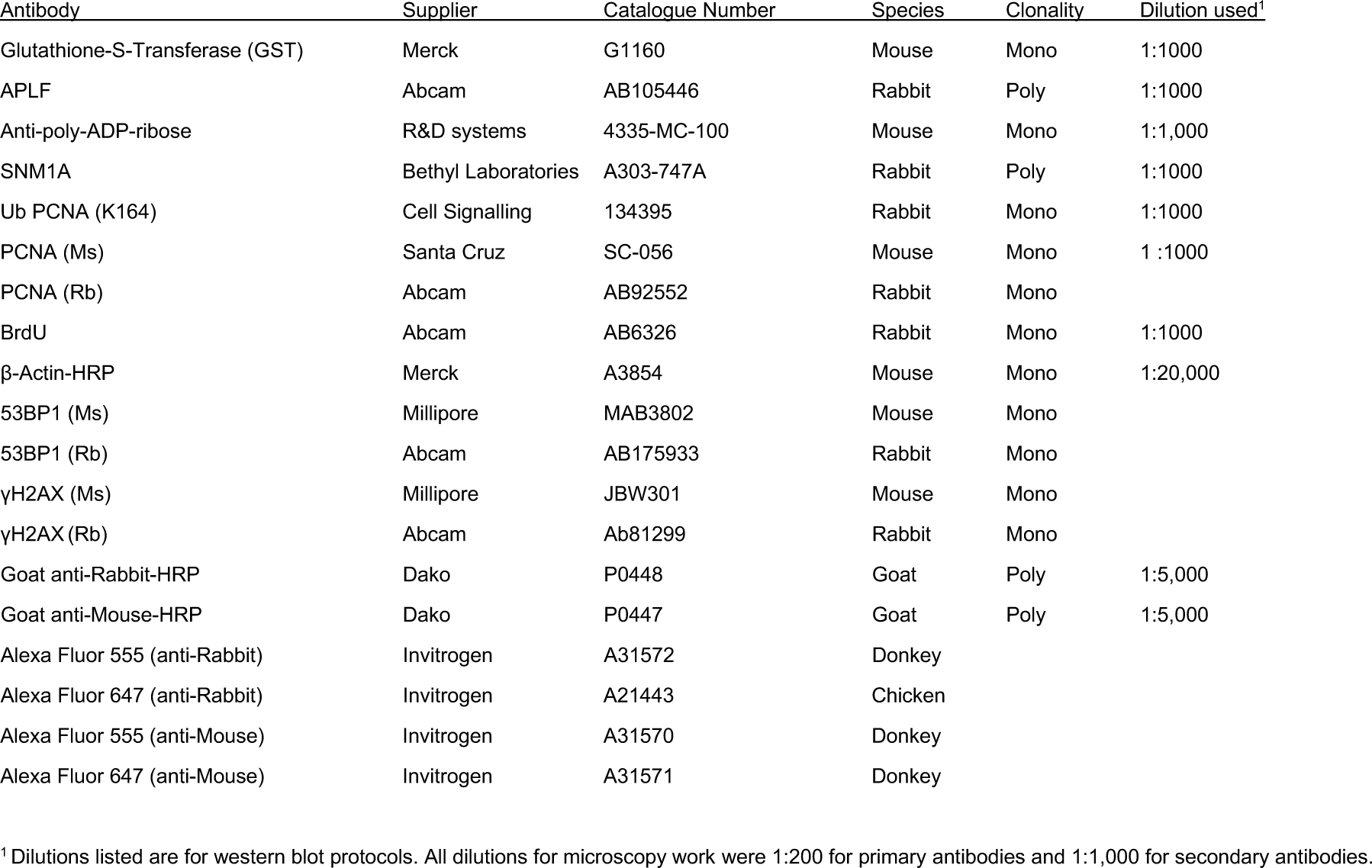

List of antibodies used in this study:
| Antibody | Supplier | Catalogue Number | Species | Clonality | Dilution used<sup>1</sup> |
| --- | --- | --- | --- | --- | --- |
| Glutathione-S-Transferase (GST) | Merck | G1160 | Mouse | Mono | 1:1000 |
| APLF | Abcam | AB105446 | Rabbit | Poly | 1:1000 |
| Anti-poly-ADP-ribose | R&D systems | 4335-MC-100 | Mouse | Mono | 1:1,000 |
| SNM1A | Bethyl Laboratories | A303-747A | Rabbit | Poly | 1:1000 |
| Ub PCNA (K164) | Cell Signalling | 134395 | Rabbit | Mono | 1:1000 |
| PCNA (Ms) | Santa Cruz | SC-056 | Mouse | Mono | 1:1000 |
| PCNA (Rb) | Abcam | AB92552 | Rabbit | Mono |  |
| BrdU | Abcam | AB6326 | Rabbit | Mono | 1:1000 |
| β-Actin-HRP | Merck | A3854 | Mouse | Mono | 1:20,000 |
| 53BP1 (Ms) | Millipore | MAB3802 | Mouse | Mono |  |
| 53BP1 (Rb) | Abcam | AB175933 | Rabbit | Mono |  |
| γH2AX (Ms) | Millipore | JBW301 | Mouse | Mono | |
| γH2AX (Rb) | Abcam | Ab81299 | Rabbit | Mono | |
| Goat anti-Rabbit-HRP | Dako | P0448 | Goat | Poly | 1:5,000 |
| Goat anti-Mouse-HRP | Dako | P0447 | Goat | Poly | 1:5,000 |
| Alexa Fluor 555 (anti-Rabbit) | Invitrogen | A31572 | Donkey |  |  |
| Alexa Fluor 647 (anti-Rabbit) | Invitrogen | A21443 | Chicken |  |  |
| Alexa Fluor 555 (anti-Mouse) | Invitrogen | A31570 | Donkey |  |  |
| Alexa Fluor 647 (anti-Mouse) | Invitrogen | A31571 | Donkey |  |  |

<sup>1</sup> Dilutions listed are for western blot protocols. All dilutions for microscopy work were 1:200 for primary antibodies and 1:1,000 for secondary antibodies.

### Immunoblot (western blot) analysis

Immunoblot (western blot) analyses were performed as described^17^. All secondary antibodies were raised in goat (Dako, Agilent technologies) and used at 1:5000 in TBS-T (Tris-buffered saline with 0.1% Tween).

### Microscopic analysis of damage-induced subnuclear foci

To assess the number of nuclear foci following treatments, cells were plated in glass bottom dishes (as above) and allowed to attach. Following treatment, cells were fixed with 4% Formaldehyde in PBS, blocked with immunofluorescence (IF) blocking buffer (5% horse serum, 1% saponin in PBS) for 1hr before being incubation with primary antibodies at the desired concentrations in IF blocking buffer (overnight, 4°C). After washing cells three times with PBS, cells were incubated with secondary antibodies (1:500) for 2-4 hrs in IF blocking buffer, washed a further 3 times with PBS then stained with Hoechst 333258 (1µg/mL in PBS) for 30 minutes before a further three washes (all washes and incubations were performed at room temperature unless otherwise stated).

Confocal images were obtained with a Plan APO 63X 1.40NA oil immersion objective, a pinhole setting of 1 AU, bandpass emission settings of 410-468 nm for Hoechst, 490-544 nm for EGFP, 579-624 nm for RFP or Alexa 568, and 633-695 nm for Alexa 647, a projected pixel dimension of around 110 nm x 110 nm, a pixel dwell time of 1.35 μs, and with a line averaging setting of 2. In order to ensure sufficient cell numbers (N>300), images were acquired in a tiled 5 x 5 format corresponding to an image area of about 0.65 mm x 0.65 mm. Images were imported into ImageJ and foci were counted using a macro script adjusting for staining levels between experiments.

Images were processed and analysed using imageJ/Fiji macro scripts^60^. Multi-channel images were split into specific fluorescent channels and then the DAPI channel was segmented to extract nuclear regions with additional splitting of cellular instances using a watershed filter. The green (EGFP), red (Alexa Fluor 568) and far red (Alexa Fluor 647) channels were filtered with a mild (0.8 sigma) Gaussian blur kernel and then regions corresponding to cells identified through correspondence with the segmentation. From these processed channels, foci were detected in each cell using the Fiji “Find Maxima” algorithm within the two channels, to create two sets of points for each cell. Each point was analysed for size and also its distance to its nearest neighbour in the comparative channel. Foci were filtered based on their size and distance, and then counted and also the intensity measured in one or more channels for each focus, depending on the experiment, and statistics for each cell established.

The macro script used can be found at: https://gist.github.com/dwaithe/ca774de63fdaae5a8e65a1d0059d61dc

### Isolation of GST-SNM1A peptides

To establish the role of binding of SNM1A and mutants with PAR chains, PCNA and ubiquitinated PCNA (PCNA^ub^) we generated GST-SNM1A peptide constructs. Consideration was given to the surrounding sequence when designing these peptides - a multiple sequence alignment was used to define the borders of the UBZ, PBZ domains and PIP box (Suppl. Fig. 5). Extra residues both upstream and downstream were included being 2+2 in the UBZ/PBZ peptide and 10+12 in the PIP box peptide.

The peptide sequences used were (the UBZ domain was mutated at C125F (highlighted in turquoise). The PBZ domain was mutated to give C155A and C161A (highlighted in yellow)):

RPVYDGYCPNCQMPFSSLIGQTPRWHVFECLDSPPRSETECPDGLLCTSTIPFHYKRYTHFLLAQSRAG.

The PIP box peptide was mutated to give Y562A and F563A (highlighted in red):

ARHPSTKVMKQMDIGVYFGLPPKRKEEKLL:

Sequences encoding for these peptides were cloned into the pET-41b vector (Novagen) using the restriction sites MfeI and SalI using standard methods and verified by sequencing.

Plasmids were transformed into competent BL21 *E . coli* cells by heat-shock. Cultures (250 mL) were grown in 2 x YT media at 37°C until OD 0.6 was reached at which point the incubation temperature was reduced to 16°C and protein expression was induced with IPTG (0.5 mM). Following 12-14 h incubation cells were pelleted in 50 mL volumes and snap frozen before being stored at −80°C.

### Preparation of recombinant PCNA and PCNA^ub^

Human PCNA was produced with a His6-tag in *E. coli* BL21 and purified by immobilized metal affinity chromatography (IMAC) followed by gel filtration using a Superdex 200 10/300 GL column (Cytiva) and 50 mM HEPES, 200 mM NaCl, 1 mM DTT, 10% glycerol, pH 7.5 as running buffer. Monoubiquitylation of human PCNA was performed in vitro as described before^61^, using a mutant UbcH5c (S22R) for conjugation. Following the reaction, PCNA^ub^ was purified essentially as described^61^ by anion exchange chromatography followed by gel filtration. For the anion exchange chromatography, a Mono Q 5/50 GL 1 mL column (Cytiva) was used. Gel filtration was performed as above.

### PCNA and PCNA^ub^ binding assay

GST-peptides were isolated by resuspending a 50 mL *E. coli* pellet in 7.5 mL of TBS- N (Tris buffered saline with 0.1% NP-40) and supplemented with 1 x protease inhibitor cocktail tablet (Merck). Cells were sonicated for four rounds of one minute on, one minute off, on ice, to release the cellular contents. Lysates were centrifuged (20,000 x g, 30 minutes at 4°C); the cleared lysates were added to pre-blocked glutathione-agarose beads (ThermoScientific, 400 μL of beads per sample, blocked with 5% FCS in PBSA for 1 hour). Binding was performed at 4°C for 2 hours. Peptide-bound beads were washed with varied concentrations of PBS (salt at 250 mM, 500 mM, 1 M, 500 mM, 250 mM) before being resuspended in standard PBS. Samples were then divided and binding to PCNA or PCNA^ub^ performed. Purified PCNA or PCNA^ub^ (150 ng) was added to each sample and tumbled at 4°C for 1 hour. Washes were performed as above to remove unbound PCNA/PCNA^ub^. The washed beads were resuspended in 2 x Laemmli buffer and boiled before being run on 4-12% SDS-PAGE gels and transferred to PVDF membrane for immunoblotting with anti-SNM1A, PCNA, PCNA^ub^ and GST.

### PAR Binding assay

To assess the binding of PAR (poly-ADP-ribose) chains to the ligand-binding motifs in SNM1A we utilised the GST-SNM1A PBZ/UBZ peptides described above. Native GST-SNM1A peptides were extracted from BL21 cell lysates (as described for PCNA binding assay above) and the peptides were eluted from the beads with excess glutathione. The peptides were then dot-blotted to PVDF. The membrane containing the GST-SNM1A peptides as well as BSA and a known PAR interacting purified protein (APLF1) as controls.

The membranes were blocked with 10% skimmed milk powder in PBS (10% milk) and a solution of Poly(ADP-ribose) polymers (1:1000 PAR chains in 10% milk, Trevigen) were incubated at room temperature (1 hr) to allow binding of the PAR chains to the proteins/peptides bound to the membrane. Following extensive washing (3 x 3 minutes) in TBS-T (Tris-buffered saline solution with 0.1% Tween 20), membranes were probed with an anti-poly-ADP-ribose (1:1000 in 10% milk, Merck) as well as anti- APLF and anti-GST as loading controls.

### Nuclease assay; substrate preparation and assay conditions

Full-length SNM1A and its variants were purified as described^17^. Human EXO1b was purified as described^62^ by the Oxford Protein Production Facility, using constructs kindly provided by Paul Modrich. Nuclease assays were performed as described^43^. Oligonucleotide substrates including those containing terminal and internal 8-oxoguanine, thymine glycol, and hypoxanthine lesions were from Eurofins, MWG Operon, Germany. The sequences and features of the oligonucleotides used are described in detail elsewhere^21^.

Substrates were prepared as follows: 10 pmol of DNA oligonucleotide (Eurofins MWG Operon, Germany) were 3’-end labelled with 3.3 pmol of α-32P-dATP (Perkin Elmer) using terminal deoxynucleotidyl transferase (TdT, 20 U; ThermoFisher Scientific), incubated together at 37°C for 1 h. Unincorporated nucleotides were removed using a P6 Micro Bio-Spin chromatography column (BioRad). For preparation of double-stranded substrates, radiolabelled single-strand oligonucleotides were annealed with the appropriate unlabelled oligonucleotides (1:1.5 molar ratio of labelled to unlabelled oligonucleotide) by heating to 95°C for 5 min, then slowly cooling to room temperature in annealing buffer (10 mM Tris–HCl; pH 7.5, 100 mM NaCl, 0.1 mM EDTA).

Nuclease assays were performed in 10 μl final volume mixtures containing 20 mM HEPES-KOH, pH 7.5, 50 mM KCl, 10 mM MgCl_2_, 0.05% Triton X-100, 5% glycerol and ∼20 nM of full-length SNM1A, 1 nM ΔSNM1A or 50 nM EXO1. Reactions were initiated by addition of DNA (10 or 100 nM), with incubation at 37°C for the indicated time period. Reactions were terminated by adding 10 μl stop solution (95% formamide, 10 mM EDTA, 0.25% xylene cyanol, 0.25% bromophenol blue) (95°C for 3 min). To examine the effects of PAR chains on SNM1A activity, 5 minutes pre-incubation with PAR (+=100nM, ++=1000nM) or an excess of PARP1 (200 nM) and NAD+ (200 µM) were performed.

Reaction products were analysed using 20% denaturing polyacrylamide gel electrophoresis (40% solution of 19:1 acrylamide:bis-acrylamide, BioRad) containing 7 M urea (Sigma Aldrich) in 1× TBE (Tris-borate EDTA) buffer at 700 V for 75 min. Gels were fixed for 40 min in a 50% (v/v) methanol, 10% (v/v) acetic acid solution, and vacuum-dried at 80°C for 2 h. Gels were imaged with using phosphorimager screen and scanned using a Typhoon 9500 phosphorimager (GE).

### SNM1A processing of Zeocin induced DNA damage *in vitro* assays

To induce Zeocin damage, DNA plasmids (3.6 μg of pG46) were treated with Zeocin in 1 x reaction buffer^63^ (12.5 mM Tris–HCl pH 8.0, 300 mM sucrose, 0.0188% Triton X-100, 1.25 mM EDTA, 5 mM MgCl_2_ with freshly added 7.5 mM β-mercaptoethanol, 1% heat-inactivated BSA and 100 mM ferrous ammonium sulphate) in 50 μL volumes for 20 minutes at 37°C. PCR clean-up columns (Qiagen) were used as per manufacturer’s instructions to clean up the reacted plasmid DNA and eluted in a 50 μL volume in buffer EB (Qiagen).

The ability of SNM1A to digest Zeocin-induced damaged DNA was measured using a Hind III linearised plasmid (a preferred substrate for SNM1A) by incubating with varying concentrations of SNM1A for 1 hour at 37°C. Following digestion, the samples (3:1) were diluted in stop buffer (95 % v/v Formamide, 10 mM EDTA, 0.25% v/v Bromophenyl blue) and resolved in 1% agarose gels in 1 x TAE buffer and visualised with ethidium bromide.

### Homologous Recombination Repair (HR) and Non-Homologous End-Joining repair (NHEJ) reporter assays

HR and NHEJ repair assays were performed as described^64, 65^. Plasmids pCMV-SceI and pDR-GFP (a kind gift from Valentine Macaulay) were used to measure HR and pimEJ5GFP (Addgene # 44026) and pCMV-SceI were used to measure NHEJ^65^. Plasmids (2.6 μg of each) were transfected into 5 x 10^5^ cells (293FT, 293FT cells treated with siRNA to BRCA2, XRCC4^-^ or SNM1A^-^ 293FT cells) in 2 mL in wells of a 6-well plate, as described above. After 24 hrs cells were harvested and resuspended in fresh media containing no phenol red and analysed on an Attune flow cytometer (ThermoFisher Scientific) to assess GFP positive cells as a measure of repair in each pathway.

---

**Suppl. Fig. 1.**
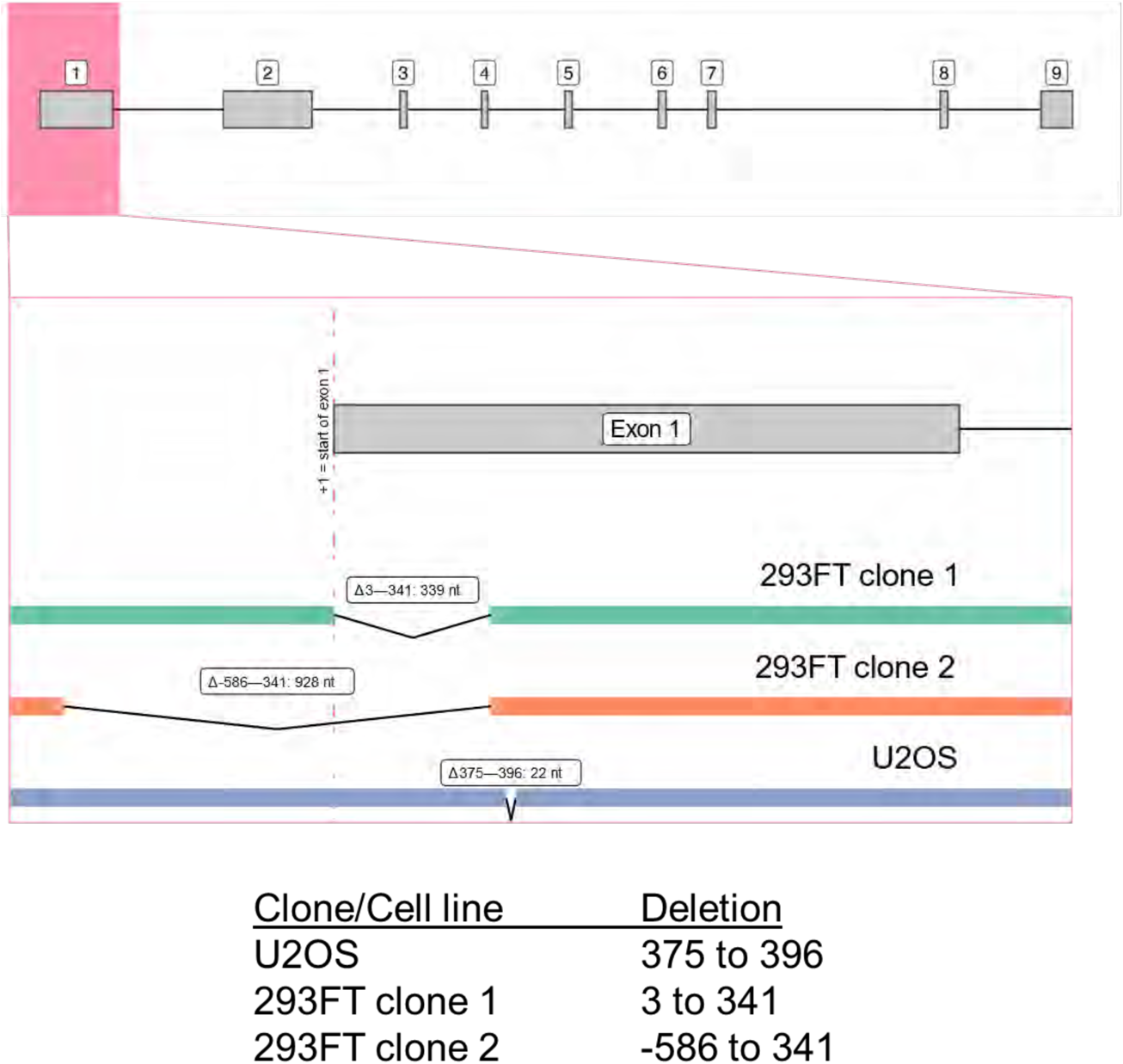
Description of deletions introduced in SNM1A gene in the cells used in this study. Regions of SNM1A deleted in disrupted cell lines. Upper panel; schematic of exon-intron structure of SNM1A. Lower panels and table; expanded views and coordinates of the nucleotide deletions within exon 1 of the gene in the U2OS-derived disruptants and in the two 293FT clones employed.

**Suppl. Fig. 2.**
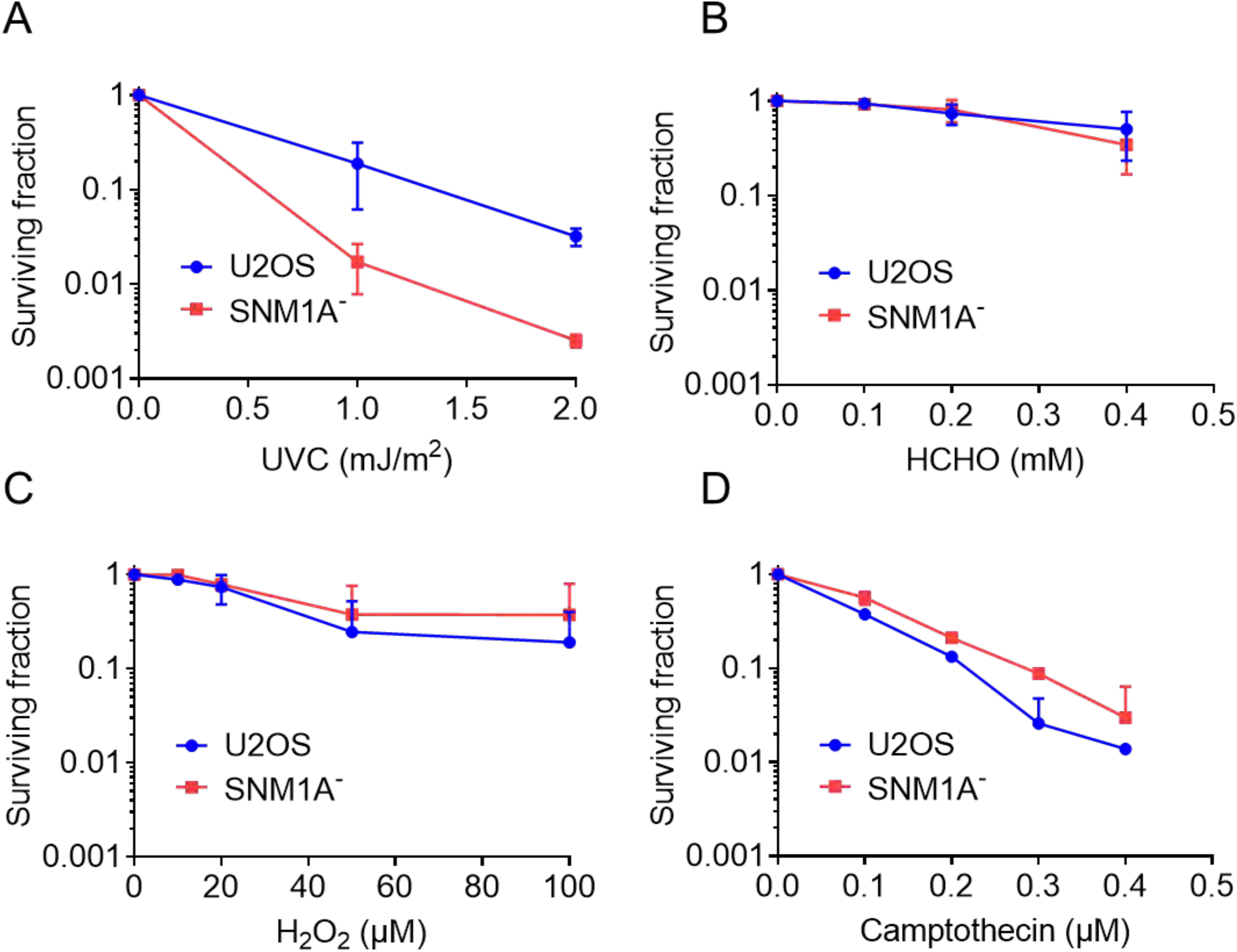
Clonogenic survival assays in U2OS and SNM1A^-^ cells. Clonogenic survival assays were performed on U2OS and SNM1A^-^ cells to treatments of UVC (254 nm) (**A.**) Formaldehyde (HCHO, **B.**), hydrogen peroxide (H_2_O_2_, **C.**) and Camptothecin (**D.**). Treated cells were allowed to grow for 12 days before being stained with Coomassie Brilliant Blue R250. Colonies were counted and the data plotted for at least 3 biological repeats containing duplicate plates per dose and normalised to the control (untreated). Error is the standard error of the mean (SEM).

**Suppl. Fig. 3.**
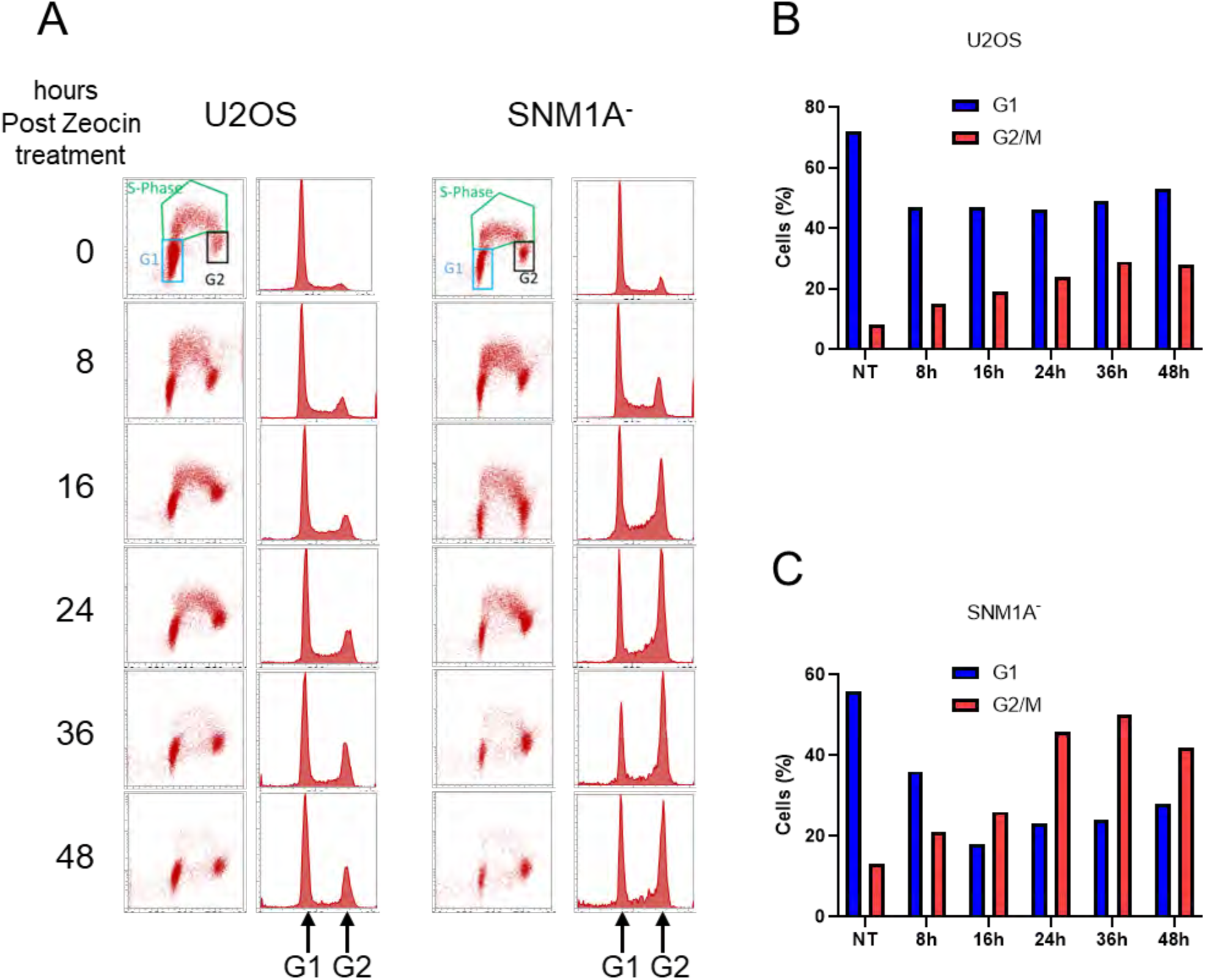
SNM1A^-^ cells accumulate in G2/M phase of the cell cycle in response to Zeocin treatment. Complete set of flow cytometry data as summarised in Fig. 1J. **A.** U2OS and SNM1A^-^ cells were treated with Zeocin (0.1 mg/mL continuously) and the cell cycle distribution analysed by BrdU incorporation. The accumulation of SNM1A^-^ cells in G2/M-phase of the cell cycle can be seen from 16 hrs and this population persists through to 48 hrs. To enumerate this accumulation, the acquired data was gated to indicate G0/G1- phase cells (G1), S-phase and G2/M-phase (G2) populations and the percentage of cells in G1- compared with G2-phase of the cell cycle was plotted for **B.** U2OS and **C.** SNM1A^-^ cells. The data displayed is representative of 3 biological repeats.

**Suppl. Fig. 4.**
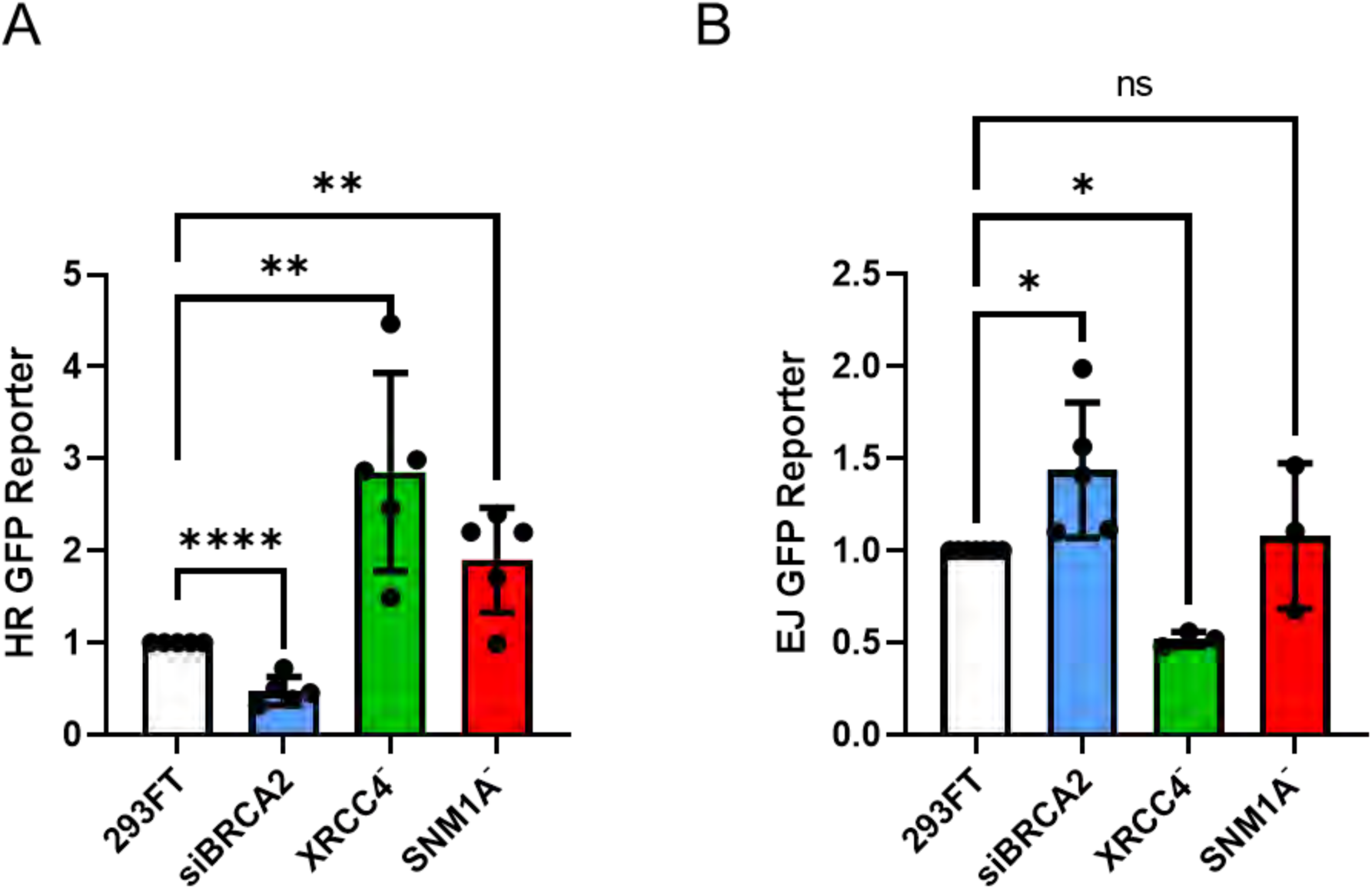
SNM1A^-^ Cells do not exhibit fundamental defects in homologous recombination or non-homologous end-joining. Wild-type cells (293FT), along with cells deficient in homologous recombination repair (HR, treated with siRNA directed against BRCA2), cells deficient in non-homologous end-joining (NHEJ, genomic deletion of XRCC4) or SNM1A^-^ cells were assayed for repair of double-strand breaks using I-SceI-reporter assays. Transfection of wild-type, siBRCA2, XRCC4^-^ and SNM1A^-^ with the HR reporter plasmid pDR-GFP **A.** or the NHEJ reporter plasmid pimEJ5GFP **B.** along with a plasmid containing the rare cutting enzyme I-SceI (pCMV-SceI). Co-transfected cells were incubated overnight, harvested, and analysed for GFP positive cells by flow cytometry. GFP positive cells for both these reporter assays indicate proficiency in repairing double-strand breaks (DSBs) induced by the co-transfected I-SceI containing plasmid. Data presented is the mean (error SEM) of three or more biological repeats. Student’s t test shows significance of the distribution where * = P≤0.05, ** = P≤0.01, *** = P≤0.001 and **** = P≤0.0001.

**Suppl. Fig. 5.**
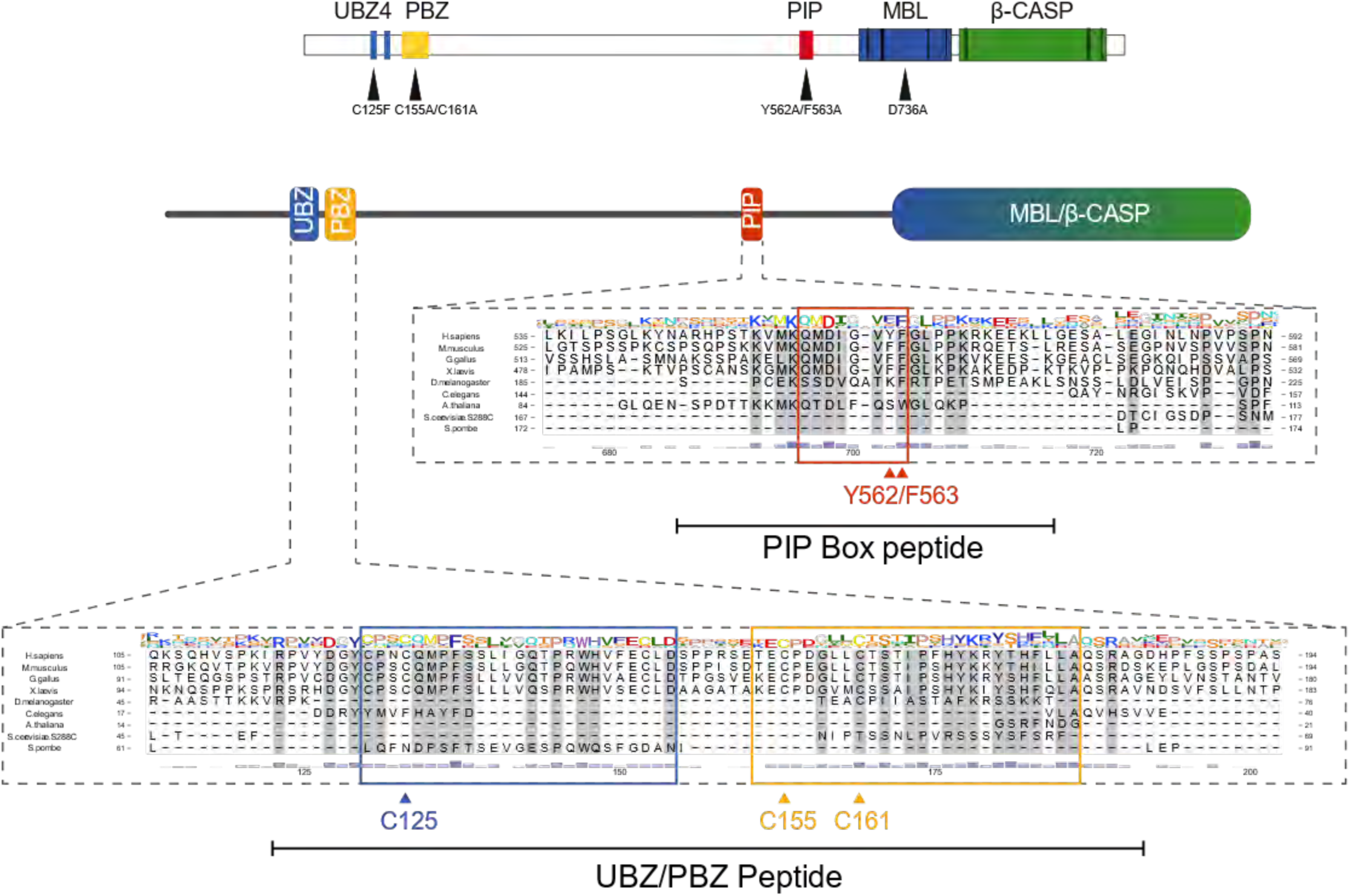
Conserved domains and motifs of SNM1A. Schematic of domains and motifs of the human SNM1A protein, with highlighted sequence alignments for the ubiquitin-binding zinc-finger 4 (UBZ), PAR-binding zinc-finger (PBZ) and PCNA-interacting peptide (PIP) box motifs. The mutated residues used in this study are highlighted, namely UBZ C125 (light blue), PBZ C155, C161 (yellow) and PIP box Y562, F563 (red), all which show strong conservation across vertebrates. The GST-SNM1A peptides used in Figure 3 to analyse the PAR, PCNA and PCNA^ub^ binding are also indicated.

**Suppl. Fig. 6.**
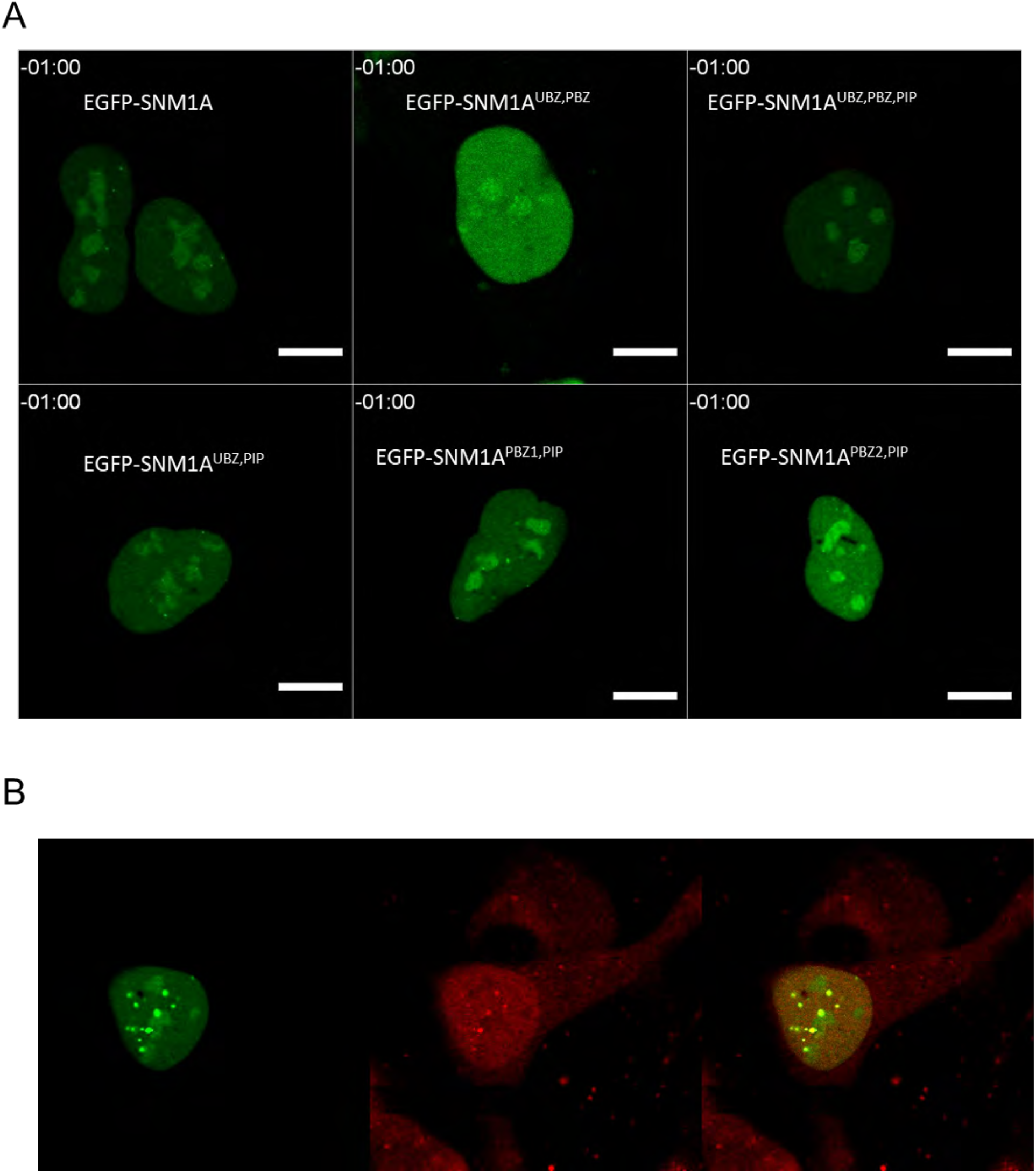
Movies of recruitment of EGFP-SNM1A its mutant forms and RFP-PCNA laser-induced DNA damage. **A.** Representative examples of movies acquired following laser induced DNA damage for wild-type EGFP-SNM1A, as well as combinations of mutants in the UBZ PBZ and PIP box. **B.** representative movie of EGFP-SNM1A and RFP-PCNA recruitment to site of laser stripe damage. Scale bars are 10 µm.

**Suppl. Fig. 7.**
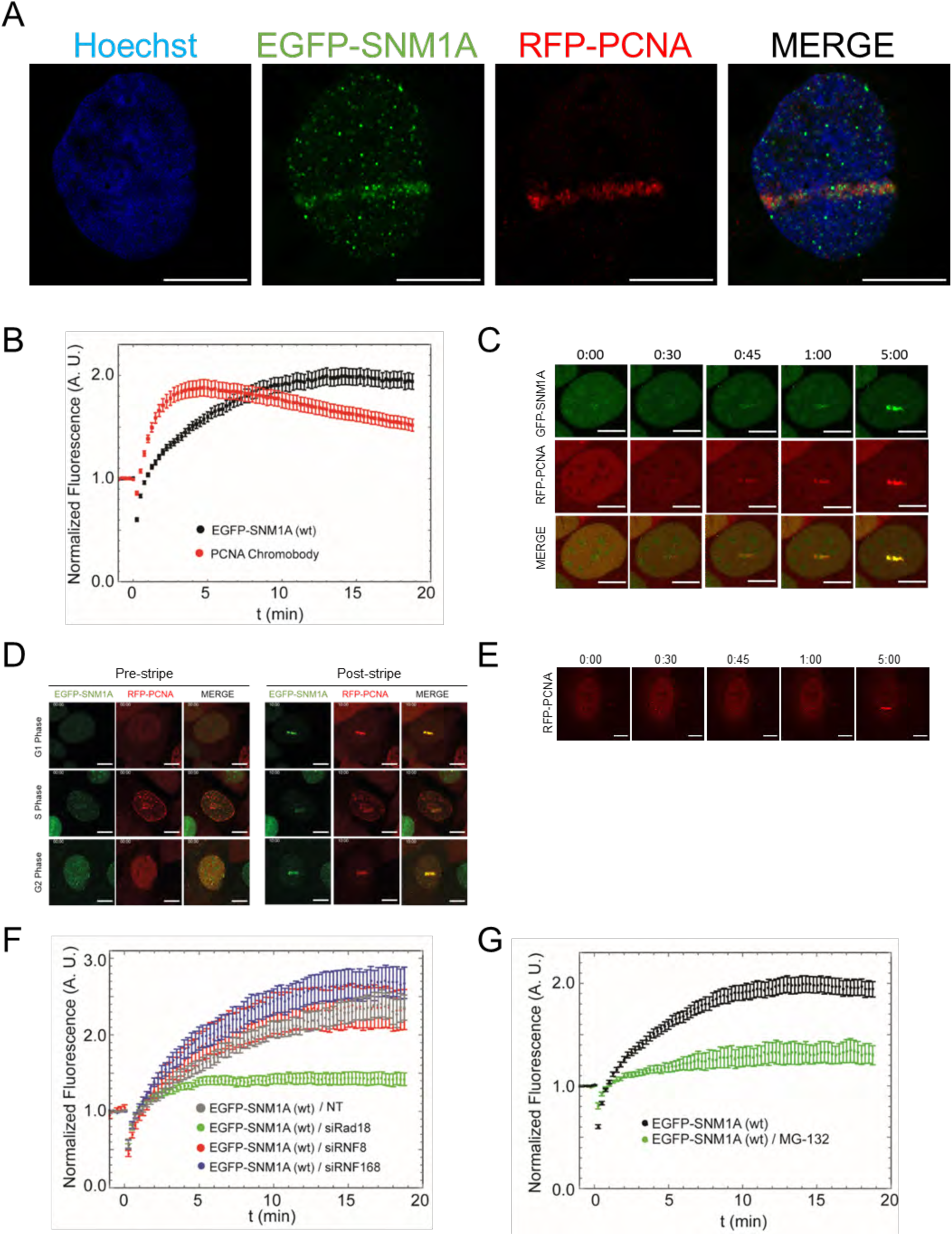
PCNA ubiquitination plays a role in SNM1A recruitment to DNA damage. **A.** Cells stably expressing EGFP-SNM1A and an RFP-PCNA Chromobody^®^ were subject to laser-induced DNA damage, where co-localisation of EGFP-SNM1A (green) and PCNA (red) to the site of laser-induced DNA damage was observed. DNA was stained with Hoechst 333258 (blue) **B.** the rate and amount of fluorescence for EGFP-SNM1A and RFP-PCNA were plotted showing that the recruitment of PCNA precedes EGFP-SNM1A. **C.** ‘snap shots’ of recruitment of EGFP-SNM1A and RFP-PCNA to laser induced damage over the first 5 minutes following laser damage and **D.** representative snapshots of recruitment of EGFP-SNM1A and RFP-PCNA in different phases of the cell cycle (note: this is an extended figure to data presented in Fig. 4E). **E.** RFP-PCNA recruitment in SNM1A^-^ cells. **F.** Effect of siRNA depletion of the major E3 ligases involved in DSB repair, Rad18, RNF8 and on the recruitment of EGFP-SNM1A to sites of laser damage. **G.** The effects of treatment with the proteosome inhibitor MG132 on EGFP-SNM1A recruitment to laser damage. All laser striping experiments represent data from at least eight cells from three biological repeats, error bars are standard error of the mean. Scale bars in A and C are 10 µm.

**Suppl. Fig. 8.**
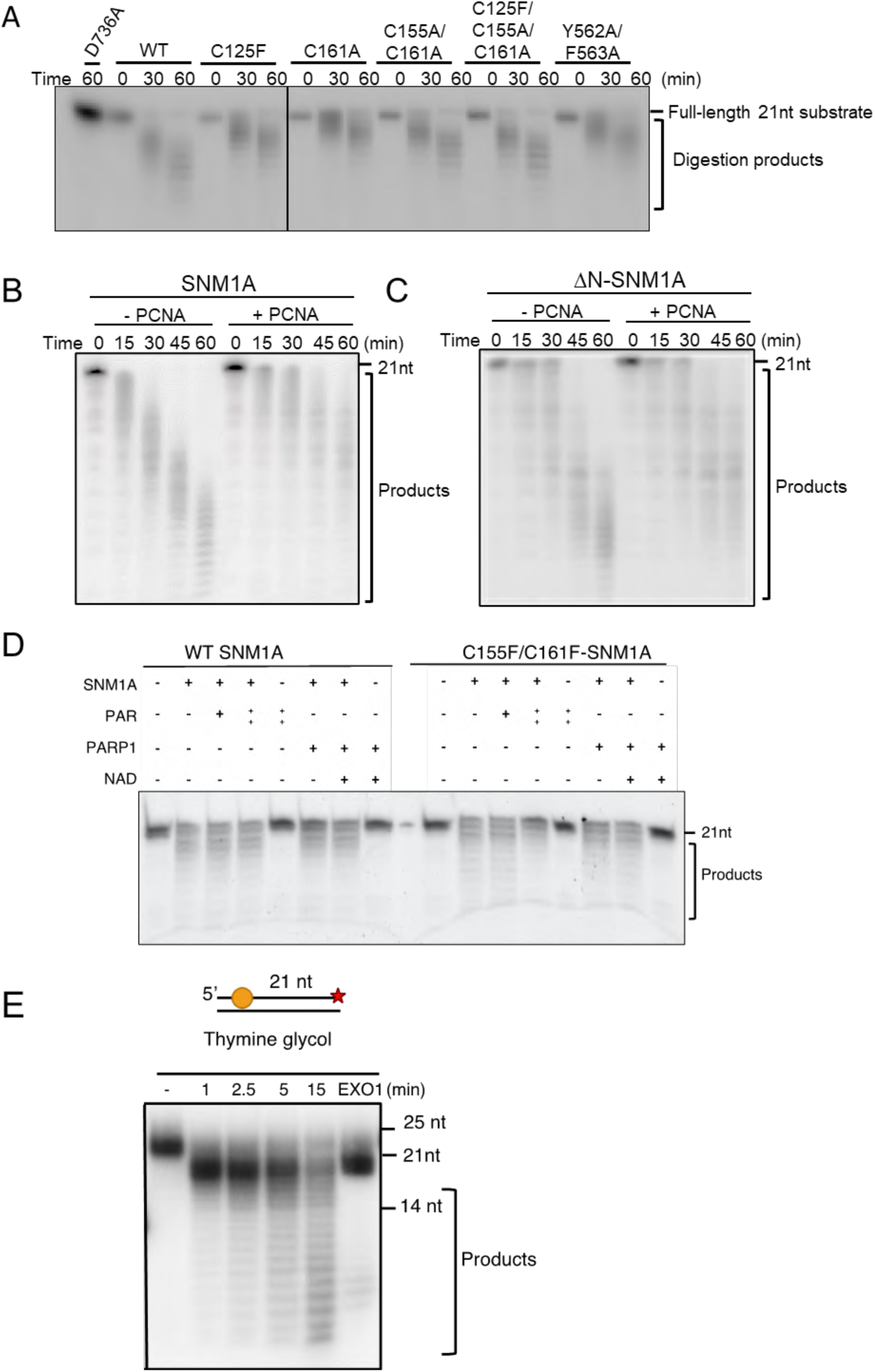
Mutations in SNM1A damage recruitment domains (UBZ, PBZ, PIP box) do not alter the enzymatic activity of SNM1A and neither PCNA nor PAR chains impact on SNM1A biochemical activity. **A.** The digestion of 21-mer ssDNA oligonucleotide by full-length SNM1A, and SNM1A bearing mutations in the UBZ (C125F), PBZ (C161A, C155A), and PIP Box (Y562A, F563A), and combinations of these. **B.** Effect of PCNA on ssDNA digestion characteristics employing full-length SNM1A, or a truncated form consisting of the MBL-β-CASP fold catalytic domain of SNM1A (ΔN−SNM1A; residues 697-1040). **C.** Effect of poly-ADP-ribose chains (PAR) directly added to the reaction mix on the digestion of a 21-mer ssDNA oligonucleotide by SNM1A. Also shown is the effect of coincubation of PARP1 with NAD^+^ to produce auto-PARylated PARP1, and the effect of this reaction on SNM1A activity. Reaction time was 30 minutes. **D. E.**

**Suppl. Fig. 9.**
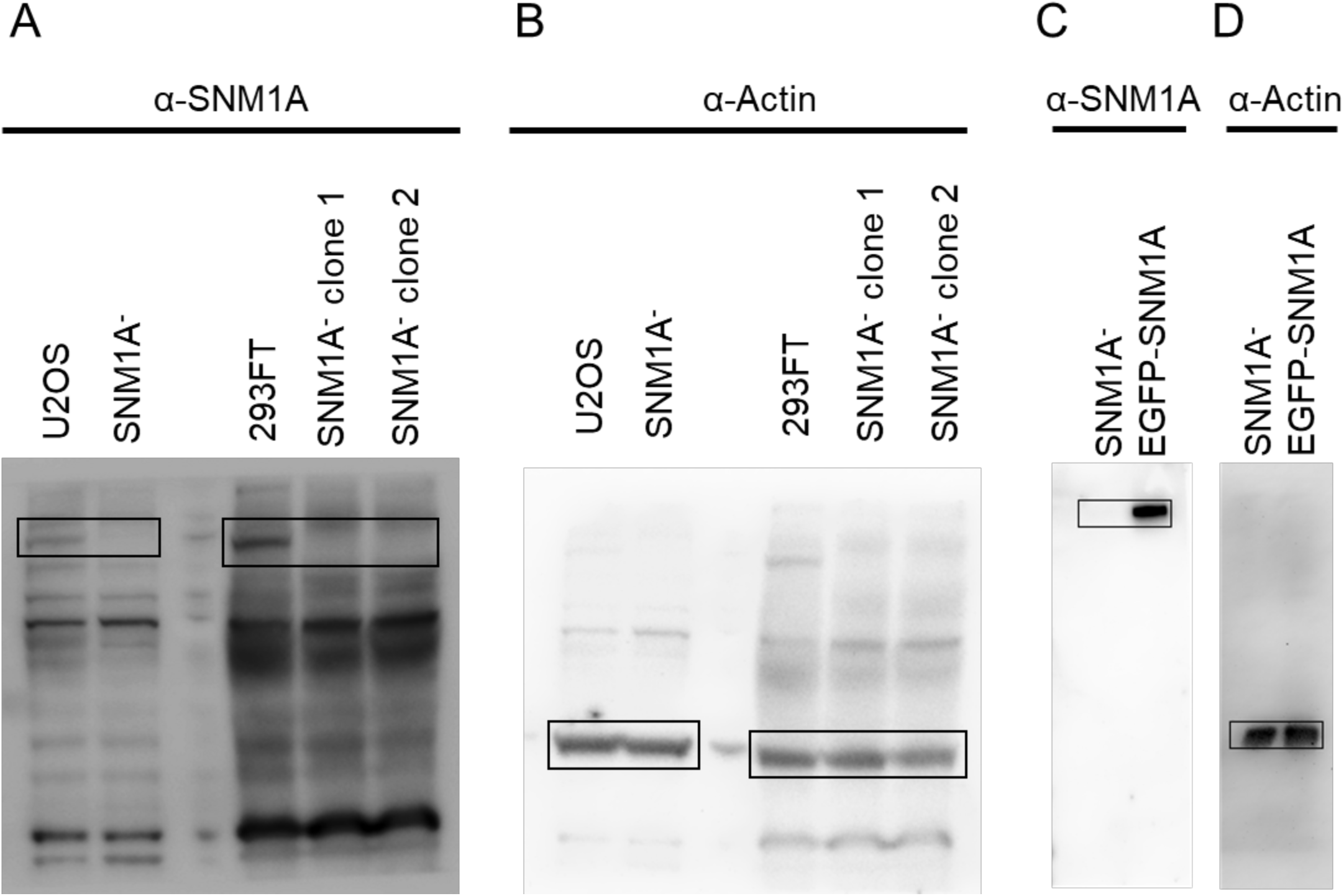
Uncropped membranes for SNM1A and GFP-SNM1A western blot analysis of protein levels in wildtype and mutant cell lines. Full blot membranes for images that appear in Figure 1A for anti-SNM1A (**A.)** and anti-actin (**B.)** in U2OS and SNM1A^-^ cells as well as 293FT and SNM1A^-^ clones1 and 2. U2OS SNM1A^-^ cells and GFP-SNM1A complemented cells probed for SNM1A (**C.)** and actin (**D.)**.

**Suppl. Fig. 10.**
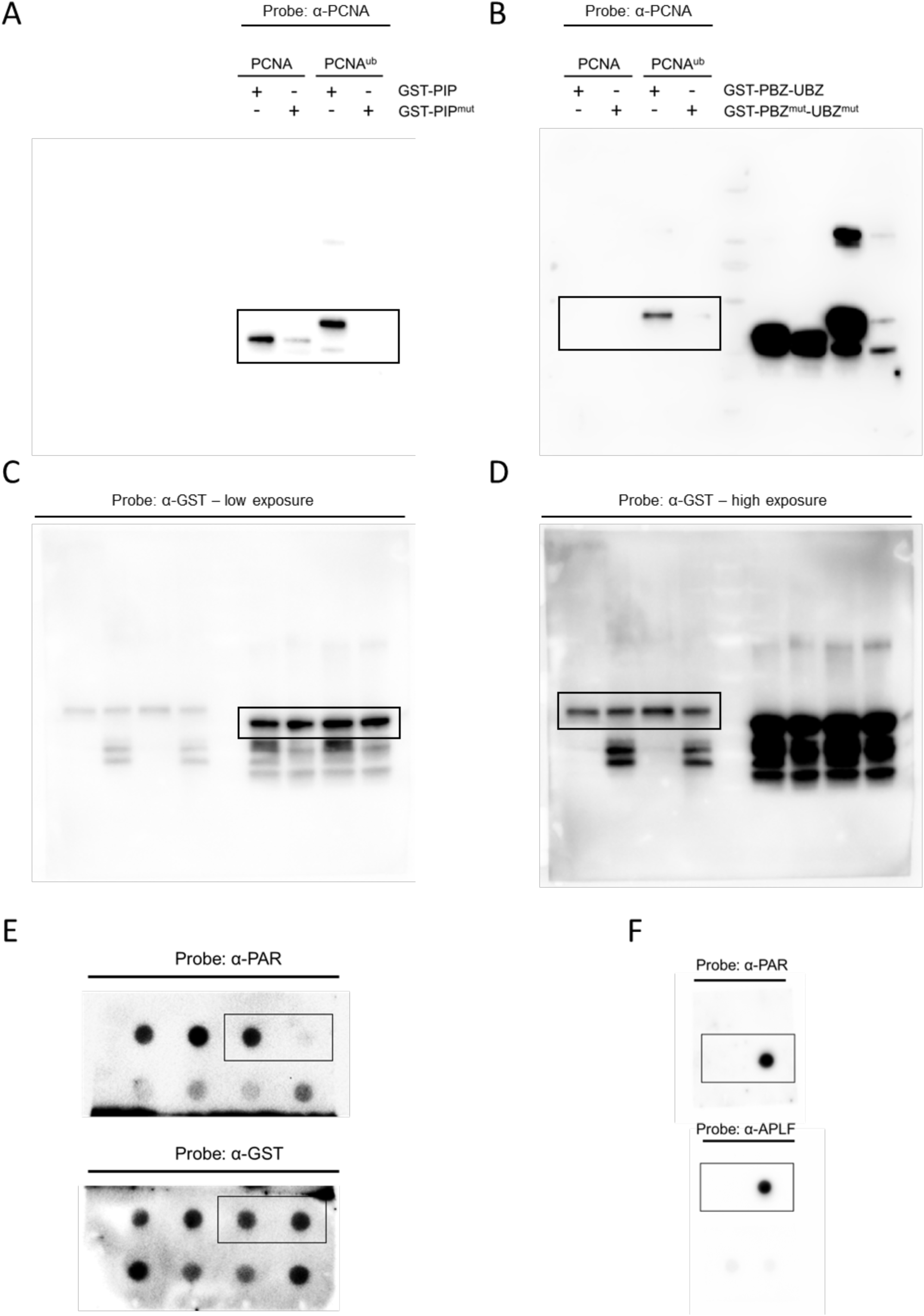
Uncropped membranes for PAR chain, PCNA and PCNA^ub^ binding to SNM1A GST peptides for PBZ, UBZ and PIP box domains. Full blot membranes of the GST-SNM1A peptide binding pull downs and dot blots from images in Figure 3 **C and D**. Probing with anti-PCNA antibodies for **A** GST-PBZ-UBZ and GST-PBZ-UBZ^mut^ peptide binding to PCNA and PCNA^ub^ (relates to Figure 3C left hand panel) and **B** GST-PIP and GST-PIP^mut^ peptide binding to PCNA and PCNA^ub^ (relates to Figure 3C right hand panel). **C.** and **D.** Probing with anti-GST antibodies for loading control for blots in **A**. and **B**. respectively. **E.** Uncropped membranes of the dot blots showing wildtype and mutant GST-UBZ-PBZ and GST-UBZ-PBZ^mut^ peptide binding to PAR chains using anti-PAR and anti-GST antibodies as a loading control. **F.** The known PAR-binding protein APLF was used as a positive control; PAR chains did not bind to BSA.

